# Ectosome uptake by glia sculpts *Caenorhabditis elegans* sensory cilia

**DOI:** 10.1101/2021.02.14.430969

**Authors:** Adrià Razzauti, Patrick Laurent

## Abstract

Cilia are sensory organelles protruding from cell surface. A tight regulation of membrane receptors traffic in and out of cilia is achieved by the action of Intraflagellar Transport (IFT). Here, we show that ectosomes bud from a subset of *C. elegans* sensory cilia. Packing and disposal of ciliary receptors in ectosomes complement their retrieval by IFT. Mutations in ciliary retrieval genes increase export of the salt sensor GCY-22 from ASER neurons by ectosomes, preventing its accumulation in ASER cilia. Ectosomes are produced from two ciliary locations: cilia tip and/or cilia base. Ectosomes budding from cilia tip are released in the environment. Ectosomes produced from the cilia base are concomitantly phagocytosed by the associated glial cells. Although ectocytosis does not require glia to occur, ectosome phagocytosis by the contacting glia contributes to maintain cilia shape and sensory function. We suggest this coordinated neuron-glia interaction is required for proper cilia function.

## Introduction

Cilia are specialized sensory compartments protruding from the cell surfaces of many cell types, including sensory neurons. Sensory cilia of the olfactory and the photoreceptors neurons concentrate the signalling components required to sense and respond to chemicals or photons, respectively. Entry and retrieval of signalling components from cilia is mediated by large Intra-Flagellar Transport (IFT) trains moving membrane proteins bidirectionally along the ciliary microtubules. IFT operates together with cargo adapters such as the IFT-A and the BBSome complexes, respectively involved in entry and removal of transmembrane proteins from cilia. Defects in cilia trafficking can alter ciliogenesis, cilia structure and composition and ultimately cilia signalling (1, 2). *C. elegans* proved to be an excellent experimental system to identify and analyse the genes required for cilia function (3). Sensory cilia integrity can be evaluated by sensory responses of the animals or by neuronal uptake of lipophilic dye (DiI) via its exposed ciliated ends (4, 5). Mutations reducing cilia length cause dye filling defective phenotypes (*Dyf*) together with sensory defects (6).

Sensory organs combining glial-like cells and ciliated sensory neurons are observed across metazoans, ranging from the invertebrate sensilla to the mammalian olfactory epithelium (7, 8). In these sensory organs, glia contributes to sensory function by releasing trophic factors, recycling neurotransmitters, controlling ion balance and pruning synapses (9–11). In *C. elegans,* most of the glial cells (46 out of 50) associate with groups of ciliated neurons to form sensory organs named sensilla, these include the inner labial, cephalic, phasmid and amphid sensilla. The amphid sensilla is the primary sensory organ of *C. elegans* allowing to sense external cues. The amphids are bilateral sensory organs, each formed by 12 ciliated neurons and 2 glial cells creating an epiderm-like continuum with the hypoderm. Two glial cells, called AMphid sheath (AMsh) and AMphid socket (AMso) limit together a matrix-filled pore opened to the external environment and housing the Nerve Receptive Endings (NREs) of 8 of the 12 amphid neurons. The NRE of the other four amphid neurons (AWA, AWB, AWC, AFD) are fully embedded within AMsh (Fig 1A, 1B).

**Figure 1:**
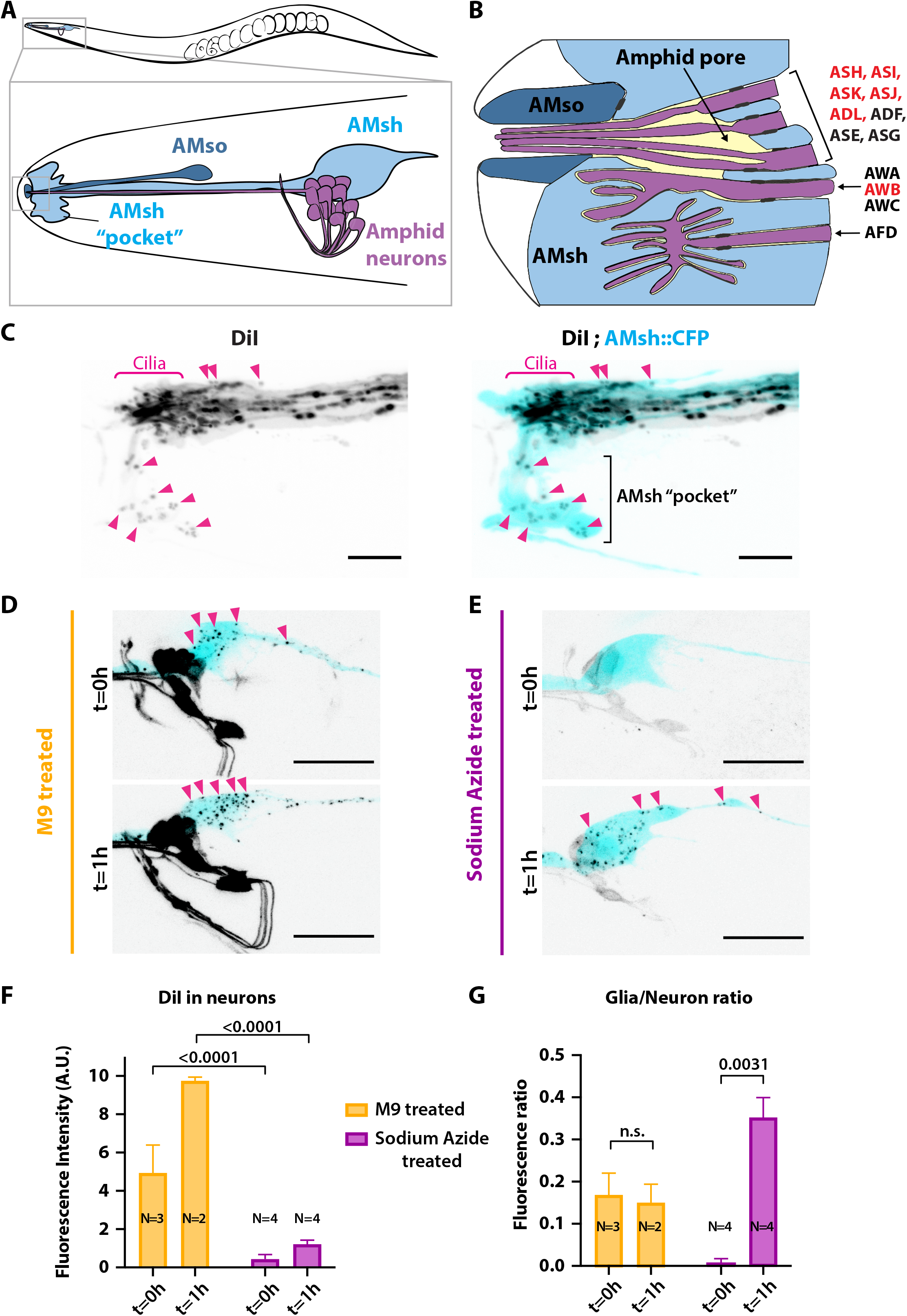
Ciliated amphid neurons transfer DiI-stained membrane to the ensheating glia in an ATP dependent manner within minutes. A) Anatomical organization of the amphid sensilla in C. elegans (Top). Head close-up scheme shows AMsh and AMsh “pocket” (light blue), AMso (dark blue) and the amphid neurons (magenta). B) Schematic depicting the NRE of the 12 amphid neurons. Tight junctions between neurons and glia, or between AMsh and AMso are depicted as dark grey discs between cells. The red labeled neurons are the DiI-stained subset. C) Maximum intensity projection of the DiI stained neurons in a strain expressing CFP in AMsh glia. Neuronally-derived vesicles containing DiI can be observed within the AMsh cytoplasm and in the AMsh “pocket” (magenta arrowheads) D) DiI staining of the amphid neurons treated with M9 only at t=0 and t=1h, amphid neurons strongly stained and multiple vesicular puncta within AMsh cell body can be observed (magenta arrowheads). E) In presence of 25mM Sodium Azide, DiI staining of the amphid neurons is fainter and no vesicles could be observed within AMsh cell body at t=0. After 1hr recovery in absence of sodium azide, the AMsh staining is recovered. F) Measurements of fluorescence intensity (in arbitrary units) quantified in neuronal cell bodies. Two-way ANOVA, Sidak’s correction for multiple comparison G) Glia/Neuron fluorescent ratios. Glia fluorescence was normalized to the fluorescence intensity of neurons. AMsh normalized fluorescence is drastically reduced in presence of Sodium Azide, and increases after its removal (t=1h). Unpaired t test with Welch’s correction. Scale bars: 5 μm in C, 20 μm in D, E.

Extracellular Vesicles (EVs) are membrane-limited vesicles released by cells. EVs hold exciting significance for biology, pathology, diagnostics and therapeutics (12). EVs are heterogeneous vesicles, including the <150 nm diameter exosomes derived from multivesicular bodies (MVBs) and the usually larger (100 nm to 1 μm) ectosomes formed via outward budding of the plasma membrane (12, 13). The capability for cilia to release ectosomes into the extracellular space has been described in a variety of organisms including *Chlamydomonas* (14–16), *C. elegans* (17), and mammals (18). Analysis of mammalian and nematode EVs content has detected several proteins that were originally enriched in the cilia, in particular polycystic kidney disease (PKD) protein polycystin-2/PKD2 (19).

In *C. elegans*, the polycystin-2 ortholog PKD-2 localises to the cilia of a subset of male ciliated sensory neurons (20). These male ciliated neurons were shown to release PKD-2-containing EVs into the external environment, where they contribute to inter-individual communication (17, 21). Electron microscopy of the male cephalic sensilla revealed EVs accumulating in the lumen surroundings of the cilia base of male cephalic neurons (CEM neurons), hinting for a release at the ciliary base (17). Different evidences suggest these male EVs correspond to ectosomes shed from the plasma membrane of the males ciliated neurons rather than exosomes derived from Multivesicular Bodies (MVBs). First, the production of ciliary EVs was not affected by mutants disrupting MVBs maturation, including mutants disturbing the ESCRT complex *stam-1, mvb-12* and *alx-1.* Second, MVBs were not observed in cilia or in the distal dendrite. Finally, some omega shaped structures were observed at the cilia base of CEM neurons (17, 22).

Several questions remain unanswered: why, where and how ectosomes bud from cilia, how cargoes enter ciliary ectosomes and what is the physiological function of these ciliary ectosomes. Here, we show that most ciliated sensory neurons of *C. elegans* pack and export their sensory receptors in two types of ciliary ectosomes. Ectosomes formed at the tip of the cilia are released to the environment while ectosomes formed at the cilia base are readily phagocytosed by the surrounding glial cells. The presence of a cargo at the time and location of ectosome biogenesis facilitates its entry into the ectosome. In addition, cargoes might contribute themselves to membrane bending or to their sorting to bending membranes. Through a candidate screen approach, we found that ectocytosis is increased in mutants defective for receptor retrieval, or in a mutant maintaining import of cilia proteins to distal dendrite in the absence of a proper cilia. These results suggest ectocytosis contributes to avoid pathological accumulations of ciliary proteins. Although ectocytosis is maintained in absence of glia, the formation of basal ectosomes is coupled to their removal by phagocytic glia. This coupling is required to maintain cilia structure and sensory function.

## Results

### Ciliated neurons transfer DiI-stained membrane to its neighboring glia in ATP-dependent manner

We used the amphid sensilla as a model of anatomically connected neurons and glia. A physical extracellular matrix barrier fills the amphid pore channel, such that only the cilia reaching the top of this pore can capture DiI. A subset of amphid sensory neurons ASH, ASI, ASJ, ADL, ASK uptake the lipophilic dye DiI from their sensory cilia. As DiI passively diffuses in lipid membranes it contacts, it is thought these cilia are in direct contact with the dye within the amphid pore (6) (Figure 1A, 1B). Despite the absence of a direct contact of AMsh with the dye the AMsh glial cells embedding the cilia base of these neurons also take up DiI, resulting in a puncta-like staining pattern, in contrast to the strong homogenous staining observed in neurons. Interestingly, this staining of the AMsh glia only occurs if the neurons themselves are stained (23). Together, these observations implied that DiI is first incorporated in membranes of the amphid neurons and secondarily the DiI-stained neuronal membranes are exported to AMsh.

Whether, where and how ciliated neurons export DiI-stained membrane to the glial cell still remains unknown. To address these questions, we analysed the DiI filling dynamics in the amphid sensilla. DiI accumulation in neurons and in glia was rapid: soaking the animals in DiI for 20 minutes was enough to stain amphid and phasmid neurons as well as the amphid and phasmid sheath glia (Figure 1C and Figure 1-figure supplement 1B). High resolution images of the nose tip revealed DiI stained vesicles within AMsh cytoplasm, either next to the dye-filled cilia or further away accumulating in a region of AMsh that we hereafter named as the AMsh “pocket” (Figure 1C). Interestingly, time-lapse imaging showed these DiI stained vesicles were produced where neuron ciliated endings are located. Once released, the flow of these vesicles was directed towards AMsh pocket or towards AMsh cell body (Video 1). Finally, DiI-containing vesicles were seen to accumulate at the soma of AMsh (Figure 1D).

Sodium azide (NaN_3_) uncouples oxidative phosphorylation and rapidly depletes intracellular ATP, disturbing membrane trafficking (24). ATP-dependent membrane trafficking events are required for the fast DiI-staining of the neurons. When worms are treated with 25mM sodium azide 15 min prior and during DiI staining, the neuron and glia staining entirely disappeared (Data not shown). However, when the animals are directly soaked for 20 minutes in a solution of DiI supplemented with 25mM sodium azide, we observed a faint staining of the neurons while AMsh glia remained unstained (Figure 1E, t=0h) (see Figure 1 – Supplement 1A for experimental scheme). This observation suggests amphid neurons captured DiI but did not transfer it to AMsh in absence of ATP. After 1 hour recovery in absence of sodium azide, the staining of AMsh was retrieved, suggesting the ATP-dependent transfer of DiI restarted (Figure 1E, t=1h). When normalising the AMsh staining to the neuronal fluorescence, DiI transfer from neurons to glia was re-established after recovery from sodium azide treatment (Fig 1F, 1G).

### The tetraspanins TSP-6 and TSP-7 enter ciliary EVs captured by the surrounding glial cells

We hypothesized that ciliary EVs released from the amphid neurons mediate the DiI export to AMsh. This scenario implies that EV markers should also be exported from amphid neurons to AMsh. To our knowledge, all current EV protein markers described for *C. elegans* are proteins mostly expressed in male sensory neurons (17, 25). In order to label ciliary EVs and show their export to AMsh, we explored other potential EV markers. The tetraspanins CD63, CD9 and CD81 are commonly used as EV markers in mammals, and tetraspanin proteins are among the most enriched proteins of human EVs (26, 27). *C. elegans* bears 20 tetraspanin genes (TSP-1 to 20). Using a reciprocal best hit approach, TSP-6 and TSP-7 appeared orthologous to CD9 and CD63, respectively (Figure 2A) (28–30). As potential ciliary EVs markers for the amphid neurons, we selected TSP-6 that is strongly expressed in the ciliated neurons and TSP-7 that is broadly expressed in the nervous system but not in ciliated neurons (31, 32) (Figure 2A). However, a valid ciliary EV marker should have two properties: 1) It should be enriched in cilia, 2) It should be loaded as a cargo in EVs released from these cilia.

**Figure 2:**
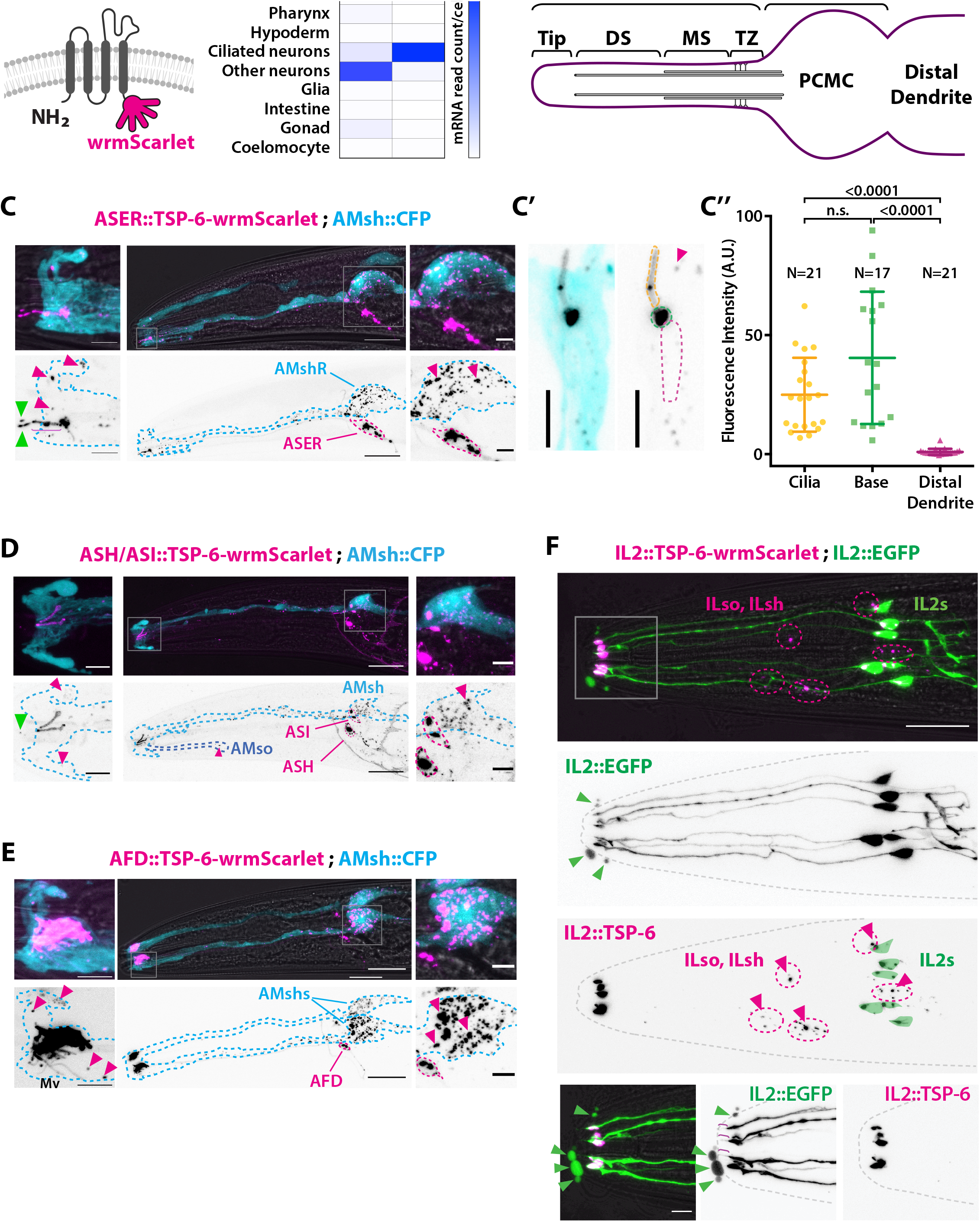
The EV marker TSP-6-wrmScarlet localizes to the cilia region, is loaded into EVs and exported to surrounding glial cells. A) TSP-6 and TSP-7 were C-terminally tagged with wrmScarlet. Expression patterns based on molecular cell profiling shows tsp-6 and tsp-7 genes are enriched in ciliated or non-ciliated neurons, respectively. B) Scheme of ciliary domains: ciliary tip, distal segment (DS), middle segment (MS), transition zone (TZ) and the periciliary membrane compartment (PCMC) located at the base of the cilia in contact with the distal dendrite. C) Expression of TSP-6-wrmScarlet in ASER driven by gcy-5 promoter. TSP-6-wrmScarlet-carrying EVs exported from ASER are observed within the cytoplasm of AMsh. Left panels show a magnification of ASER cilium region, EVs are released in the cilia pore anterior to the ASER cilia tip (green arrowhead). Right panels show EVs within AMsh cell body (magenta arrowheads). Scale bar: 20 μm middle panel and 5 μm insets. C’) TSP-6-wrmScarlet is enriched in ASER cilium: representative confocal projection showing TSP-6-wrmScarlet enrichment in PCMC and cilium C’’) Fluorescence quantification in animals expressing TSP-6-wrmScarlet in ASER neuron. Brown-Forsythe ANOVA, multiple comparisons corrected by Dunnett’s test. D) Expression of TSP-6-wrmScarlet under the sra-6 promoter (driving expression in ASH and ASI neurons). TSP-6-wrmScarlet is enriched in both ASH and ASI cilia. Left panel shows TSP-6-wrmScarlet-carrying EVs released by ASI and/or ASH in the cilia pore (green arrowhead), EVs released by ASH/ASI were also seen within the cytoplasm of AMsh surrounding ASI/ASH cilia (magenta arrowheads). Few vesicles were also observed in AMso (blue dashed outline, middle panel). Right panel shows AMsh soma with multiple EVs (magenta arrowhead). Scale bar: 20 μm middle panel and 5 μm insets. E) Expression of TSP-6-wrmScarlet in AFD neurons driven by gcy-8 promoter. TSP-6-wrmScarlet is enriched in AFD microvilli and cilia base. Left panel shows TSP-6-wrmScarlet-carrying EVs within the cytoplasm of AMsh that surrounds AFD terminals (magenta arrowheads). Right panel shows AMsh soma with multiple EVs (magenta arrowhead). Scale bar: 20 μm middle panel and 5 μm insets. F) Co-expression of TSP-6-wrmScarlet and cytoplasmic mEGFP in IL2 neurons (driven by klp-6 promoter). mEGFP can be observed within EVs that are environmentally released (green arrowhead) while TSP-6-wrmScarlet is observed on EVs located within the cytoplasm of ILsh and ILso glial cells (magenta arrowheads). Theoretical position of ILsh and ILso was outlined (magenta dashed circles), IL2 neurons position was drawn with green filled outlines. Scale bar: 20 μm top panels and 5 μm insets.

First, to answer if TSP-6 and TSP-7 were enriched in the cilia, we developed TSP-6-wrmScarlet and TSP-7-wrmScarlet fluorescent fusion proteins and generated transgenic animals expressing these constructs in specific neurons or subsets of neurons. Each neuron or group of neurons was chosen for the positioning of its Nerve Receptive Endings (NRE) with respect to their supporting glia. In each of the neurons we examined -including the ASER, AFD, ASH, ASI, ADL, ADF, AWA and IL2 neurons-TSP-6-wrmScarlet and/or TSP-7-wrmScarlet appeared enriched at the apical NREs (Figure 2 and Figure 2 - figure supplements 1 and 2). The canonical sensory cilium is subdivided in various sub-ciliary domains (Figure 2B). Quantification showed that TSP-6-wrmScarlet was enriched at the ASER cilium and PCMC compared to the distal dendrite (Figure 2C’-C’’). Similarly, using the dynein light intermediate chain XBX-1 marker to label the axoneme and PCMC, we saw that XBX-1-mEGFP largely overlapped with TSP-7-wrmScarlet location in ASH, ADL, ADF cilia, further supporting its enrichment in cilia and cilia base (Figure 2- figure supplement 1B).

Second, if TSP-6-wrmScarlet and TSP-7-wrmScarlet were loaded in EVs released by amphid neuron cilia, we should observe these markers in EVs located either in the amphid pore or exported to /captured by AMsh, as we observed for DiI. Indeed, when expressed from a subset of amphid neurons, these markers were exported from their cilia to EVs located in the amphid pore and/or in AMsh cytoplasm. Within AMsh, TSP-6-wrmScarlet and TSP-7-wrmScarlet were always observed as intracellular vesicles (Figure 2C-F and Figure 2 - figure supplement 1A, B’, C’ and Figure 2 - figure supplement 2B). However, because these vesicles originate from the neurons, we will call them EVs, something that is justified later. When expressed in the right ASE neuron (ASER), TSP-6-wrmScarlet and TSP-7-wrmScarlet were exported to fluorescent EVs located only in the right AMsh cell (AMshR) (Figure 2C and Figure 2 - figure supplement 1A). We did not observe EVs in AMso glial cells nor in the contralateral AMshL, suggesting that EV export occurs from the ASER cilum base, the only place where ASER directly contacts AMshR. Accordingly, we observed TSP-6-wrmScarlet-carrying EVs in close proximity to the ASER cilium base as well as in the AMsh pocket (Figure 2 C, insets, magenta arrowheads). We also observed fluorescent EVs in the amphid pore suggesting that these can also be apically released from ASER cilium into the amphid pore (Figure 2 C, insets, green arrowheads). Similarly, we observed export of TSP-6-wrmScarlet-carrying EVs from the bilateral ASH and ASI neurons to the amphid pore and to both AMsh cells (and rarely in AMso glia) (Figure 2D). We observed export of TSP-7-wrmScarlet-carrying EVs from the bilateral ASH, ADL, ADF, AWA neurons to both AMsh cells (Figure 2 - Figure Supplement 1B’). Finally, we assayed the AFD neuron as its NREs consist of many microvilli and a single vestigial cilium all embedded within AMsh cytoplasm. When expressed in AFD neurons, TSP-6-wrmScarlet and TSP-7-wrmScarlet were observed at the surface of microvilli and AFD base (Figure 2E - Figure 2 - figure supplement 2B). High-resolution imaging allowed us to see TSP-7-wrmScarlet in a vesicular compartment within AFD base, possibly corresponding to a recycling or trafficking compartment (Figure 2- figure supplement 2A). Similar export properties were observed for AFD neurons: EVs carrying TSP-6-wrmScarlet and TSP-7-wrmScarlet were exported from bilateral AFD neurons to both AMsh glial cells (Figure 2E and Figure 2 - Supplement 2B). Fluorescent EVs were observed within the cytoplasm of AMsh glia, around the microvilli of AFD and also in the AMsh pocket area. Fluorescent EVs were also observed in AMsh cell body (Figure 2E and Figure 2- figure supplement 2A-B). Export of EVs carrying TSP-7-wrmScarlet from AFD neurons to AMsh was observed across all larval stages from L1 stage to adult. As the animals aged, we observed a progressive build-up of TSP-7-wrmScarlet EVs in AMsh glia (Figure 2- Supplement 2B), suggesting that this export of TSP-7-wrmScarlet EVs occurs continuously from AFD neurons to AMsh and starts at early larval stages. Once exported to AMsh, TSP-7-wrmScarlet EVs might accumulate in acidified compartments belonging to the endolysosomal pathway. Compared to wrmScarlet, GFP-variants are more sensitive to quenching and degradation when subjected to acidic pH (33). When TSP-7 was fused to mEGFP, the fusion proteins were still exported from AFD neurons to AMsh, however TSP-7-mEGFP labelled EVs appeared less numerous and less bright than for TSP-7-wrmScarlet, suggesting both fusion proteins are partly located in acidic compartments after their export from AFD to AMsh (Figure 2 – figure supplement 2C). Therefore, we show that TSP-6-wrmScarlet and TSP-7-wrmScarlet label ciliary membrane compartments. These markers can be loaded into EVs that are released from amphid neurons to amphid pore or to AMsh, where they ultimately enter in an acidic compartment.

### Ectosome biogenesis from two ciliary locations produce divergent EVs fates

The cilia of male IL2 neurons were previously described to produce and release EVs to the environment (17). Accordingly, we could also observe EV release occurring from the hermaphrodite IL2 cilia expressing different markers: TSP-6-wrmScarlet co-expressed with cytoplasmic mEGFP, TSP-7-wrmScarlet co-expressed with cytoplasmic mEGFP or cytoplasmic mCherry alone. Expression of TSP-7-wrmScarlet and TSP-6-wrmScarlet appeared enriched at the IL2s cilia base compared to cilia proper, while cytoplasmic mEGFP or mCherry were homogenously distributed in IL2s cilia (Figure 2 F and Figure 2 – figure supplement C-C’’). The EVs released outside the animals were always carrying cytoplasmic mEGFP or mCherry (Figure 2 F and Figure 2 – figure supplement 1 C’’) but were only occasionally carrying TSP-7-wrmScarlet and TSP-6-wrmScarlet. In contrast, EVs exported from IL2 neurons to their supporting glia were always carrying TSP-6-wrmScarlet and TSP-7-wrmScarlet (Figure 2F, Figure 2 - Figure Supplement 1C’). The stained glia presumably corresponded to the inner labial sheath (ILsh) and socket (ILso) glial cells, based on their location. Therefore, IL2 neurons likely release two type of EVs, EVs transferred to the glia carrying TSP-6-wrmScarlet and TSP-7-wrmScarlet and EV released outside the animals that often lack the tetraspanin markers.

To analyze the dynamic of biogenesis, release and transfer of ciliary EV, we performed live imaging of animals co-expressing the cytoplasmic mEGFP along with TSP-7-wrmScarlet/TSP-6-wrmScarlet in IL2 neurons. These animals were immobilized with 10 mM Tetramisole, an agonist of cholinergic receptors that force muscular paralysis but does not interrupt ATP-dependent processes (34). In these conditions, we observed outward budding of the plasma membrane typical of ectosomes in IL2 neurons. The events occurred from two ciliary locations: first, ectosomes varying from below 250 nm to ~2 μm diameter are formed and released from the IL2 cilia tip by a constant flow of membrane towards a cilia protrusion at IL2 cilia tip. Scission of the ectosomes from the cilia tip seemed to occur randomly, releasing ectosomes of variable size in these preparations. These distal ectosomes did not always carry the TSP-7-wrmScarlet or TSP-6-wrmScarlet markers (Video 2). TSP-7-wrmScarlet appeared sorted or not into the IL2 apical ectosomes according to its presence or absence at the cilia tip at the time of ectosome budding. A ~2 μm diameter distal ectosome was formed and released in ~150 sec suggesting a fast directional membrane flow toward the cilia tip of IL2 neurons (Video 3). Second, IL2 neurons also released ectosomes carrying TSP-7-wrmScarlet from their cilia base. These were observed as translocating from IL2 cilia base towards ILsh or ILso cytoplasm (Video 2). We also performed in vivo live recordings from animals expressing TSP-6-wrmScarlet in ASER neurons. We observed a similar flow of barely detectable basal ectosomes released from ASER cilia base, suggesting local production of ectosomes whose size are close to the light diffraction limit of conventional confocal microscopes (Video 5).

Altogether, our results suggest a model where ciliary ectosomes mediate the export of ciliary membrane containing either: DiI, membrane proteins like tetraspanins (TSP-6-wrmScarlet or TSP-7-wrmScarlet) and/or ciliary cytoplasm (including mEGFP or mCherry). Ectosome biogenesis up to scission is fast but variable ranging from few seconds to few minutes, leading to variable ectosome size. These ciliary ectosomes bud from the NRE of multiple ciliated neurons, either from the cilia base and/or from cilia tip. Markers enriched at cilia base are also more likely to enter in basal ectosomes than markers enriched in distal cilia. When budding from the cilia base, ectosomes are subsequently captured by the contacting glial cells. Within these glial cells, the ectosomes enter an acidic organelle, presumably part of the endolysosomal pathway. When ectosomes are budding from ciliary tip, these are not captured by the adjacent glial cells but instead they are environmentally released.

### Endogenous ciliary receptors are sorted to ectosomes and exported to AMsh

Previous work showed that only a subset of ciliary proteins enters the *C. elegans* male EVs (21, 25). To explore which cilia proteins can enter basal ectosomes to be exported to the embedding glial cells, we fluorescently tagged several ciliary membrane proteins endogenously expressed in AFD, ASER and ASK neurons and known to localise to their cilia. The transmembrane guanylyl cyclase receptor GCY-8 is localised to AFD receptive endings where it contributes to the thermotaxis behaviour of *C. elegans* (35). Expression of the fusion protein GCY-8-wrmScarlet in AFD showed enrichment in all the distal AFD NREs, including microvilli and cilia base. GCY-8-wrmScarlet was exported from AFD to AMsh, producing fluorescent EVs that surrounded AFD microvilli and that were ultimately trafficked to AMsh cell body (Figure 3A). We also tagged a G-Protein Coupled Receptor (GPCR) endogenously localised in AFD microvilli, the SRTX-1 receptor (36). When expressed in AFD neurons, SRTX-1-wrmScarlet was also exported from AFD terminals to EVs in AMsh glia (Figure 3B). We next tagged the SRBC-64 GPCR involved in pheromone sensing and known to localise to the cilia and cilia base of the ASK neurons (37). Expression of SRBC-64-wrmScarlet in ASK neurons localised to the cilia and cilia base, but was not exported to AMsh (Figure 3C). Finally, we tagged the transmembrane receptor guanylyl cyclase GCY-22, a specific ASER cilium-located protein involved in salt sensing (38). The GCY-22-wrmScarlet fusion protein localised in a bi-partite distribution at the cilium tip and cilium base of ASER, as previously described (39) (Figure 3D). However, overexpression of GCY-22-wrmScarlet in ASER induced an enlargement of its cilium base (Figure 3D and Figure 3 - Supplement 1A). We observed that GCY-22-wrmScarlet was also exported from the cilium base of ASER to AMsh, producing fewer but noticeably larger (~1um) fluorescent EVs in AMsh cell body (Figure 3D). In addition, GCY-22-wrmScarlet was observed enriched at the membrane of large protrusions budding from ASER cilium base (Figure 3D, orange arrowhead). Because of their large size, budding of GCY-22-wrmScarlet-carrying EVs could be resolved and their budding dynamics recorded in vivo. Multiple recorded videos revealed the buddings of ~1um diameter ectosomes, starting with the elongation of a protrusion from ASER cilia base followed by the narrowing of the tubule connecting the protrusion to the cilium base up to the scission event (Video 6). All budding events took place within 6 to 25 minutes of recording (N=3). The entire budding, neck elongation and scission of basal ectosomes containing GCY-22 happened in close contact with AMsh, hinting to a phagocytic process. As soon as scission occurred, the large vesicles moved retrogradely towards AMsh cell body. We observed that different cargoes expressed in ASER produced EVs of very different size in AMsh: GCY-22-wrmScarlet generated ~1 μm EVs while TSP-6-wrmScarlet generated <500 nm EVs (Figure 3 – figure supplement 1B), suggesting the two proteins might be sorted independently to different EVs. To explore this possibility, we co-expressed GCY-22-mEGFP and TSP-6-wrmScarlet in ASER. We observed a poor overlap of the two membrane proteins within ASER cilium (Figure 3 - figure supplement 1C). Both markers were sorted to EVs and exported by ASER into AMsh, 75% of these EVs observed in the surroundings of ASER cilium were carrying TSP-6-wrmScarlet alone, 23% of them were carrying TSP-6-wrmScarlet together with GCY-22-mEGFP and very few vesicles were observed with only GCY-22-mEGFP. Interestingly, the EVs carrying GCY-22-mEGFP and TSP-6-wrmScarlet together were slightly larger than EVs carrying TSP-6-wrmScarlet alone (Figure 3-figure supplement 1E).

**Figure 3:**
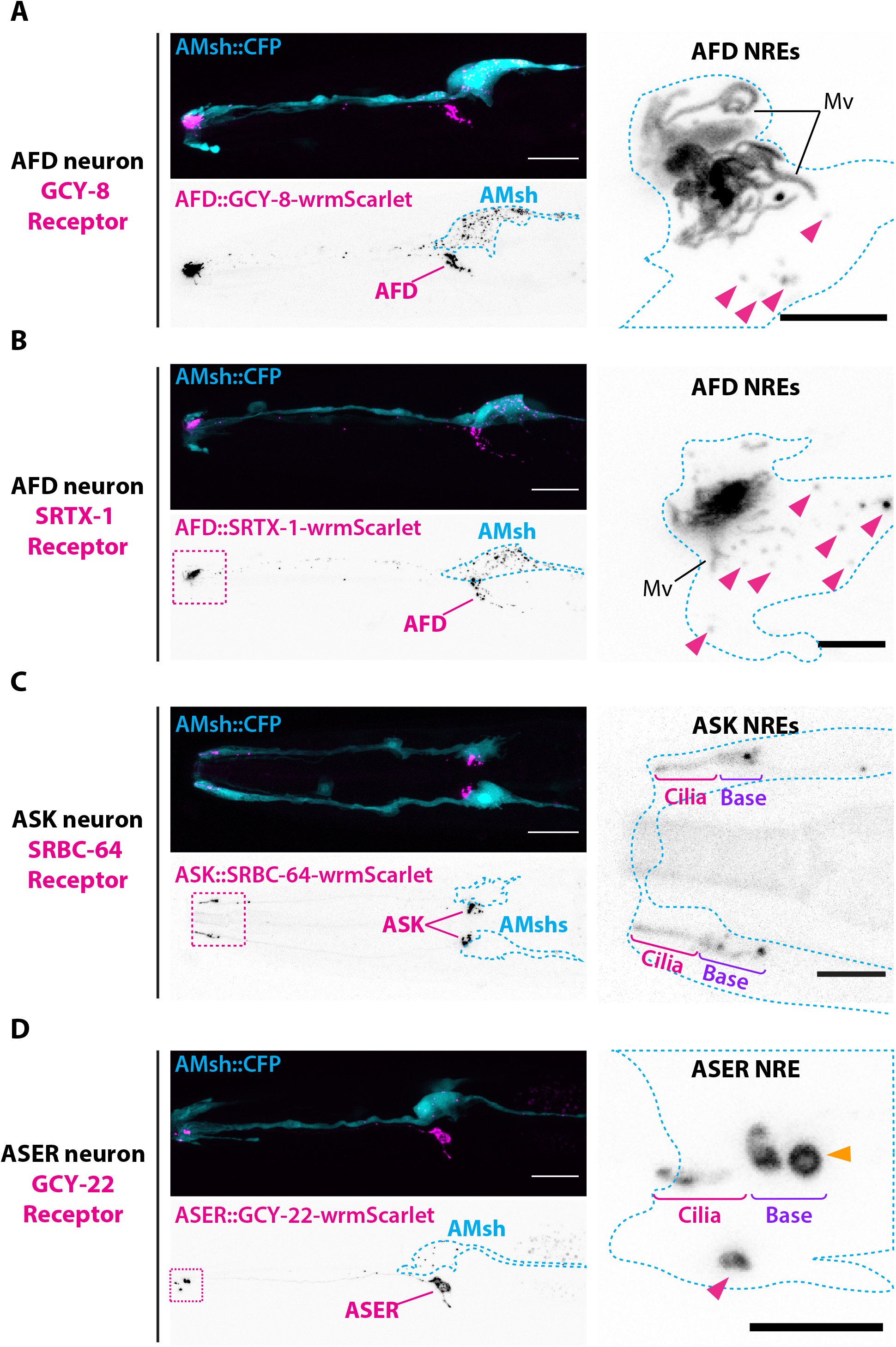
Endogenous ciliary membrane proteins are sorted to ectosomes and exported to AMsh. A) GCY-8-wrmScarlet was cell-specifically expressed in AFD (driven by gcy-8 promoter). GCY-8-wrmScarlet is enriched in AFD microvilli and AFD cilia base. GCY-8-wrmScarlet-carrying EVs were also observed within AMsh cytoplasm, in the vicinity of the AFD nerve receptive endings and in AMsh cell body (magenta arrowheads). B) SRTX-1-wrmScarlet was cell-specifically expressed in AFD (driven by gcy-8 promoter). SRTX-1-wrmScarlet is enriched in AFD microvilli and cilia base, similarly to GCY-8. Within AMsh cytoplasm, SRTX-1-wrmScarlet-carrying EVs are observed in the vicinity of the AFD neuron receptive endings and in AMsh cell body (magenta arrowheads). C) SRBC-64-wrmScarlet was cell-specifically expressed in ASK neurons (driven by srbc-64 promoter). SRBC-64-wrmScarlet is observed in the ASK cilia proper (Cilia) and cilia base (Base) but not in the cytoplasm of AMsh. D) GCY-22-wrmScarlet was cell-specifically expressed in ASER (driven by gcy-5 promoter). GCY-22-wrmScarlet is observed in ASER cilium tip and in ASER cilium base. ASER cilium base shows a rounded protrusion, that we consider as a recently excised EV (orange arrowhead). Within AMsh cytoplasm, a GCY-22-wrmScarlet-containing ectosome is located in the vicinity of the ASER cilia base. Few but large GCY-22-wrmScarlet-carying EVs are observed in AMsh cell body. Scale bar: 20 μm in head images, 5 μm in insets.

We showed that most of the ciliary membrane proteins we assessed are loaded into ciliary EVs which are readily taken up by AMsh. However, there are exception exemplified by SRBC-64-wrmScarlet, suggesting some ciliary membrane proteins do not enter EVs. We also show that ciliary proteins are not necessarily sorted together into the same ectosomes. Interestingly, we observe that ectosome composition can alter the ectosome budding properties: the cilia base of ASER expressing GCY-22-wrmScarlet is expanded and displays large protrusions that slowly bud into ectosomes. Ultimately ectosome cargo composition can alter their size: ectosomes carrying both GCY-22-mEGFP and TSP-6-wrmScarlet have an intermediate size between the ~1um of ectosomes carrying GCY-22-wrmScarlet alone and the ~400 nm ectosomes carrying TSP-6-wrmScarlet alone, suggesting cargo interaction set the EV diameter.

### Mutants defective for ciliogenesis or for cilia retrieval increase ectocytosis

Previous experiments showed that large ectosomes carrying GCY-22-wrmScarlet were formed at the cilia base of ASER and simultaneously captured by AMsh. Assuming that the number of GCY-22-wrmScarlet vesicles in AMsh cell body mostly reflects the strength of the GCY-22-wrmScarlet export by ectocytosis, these transgenic animals were crossed into mutants that potentially modulate ectocytosis. The selected set of mutants are involved at different steps crucial for the correct functioning of the cilia: ciliogenesis, cilia trafficking or cilia sensory activity. We first asked if ectocytosis required cilia to occur. In *C. elegans*, most genes involved in ciliogenesis are under the transcriptional control of the RFX-type transcription factor DAF-19 (40). *daf-19(m86)* mutant worms are completely void of ciliated structures. However, absence of cilium in *daf-19(m86)* does not impair trafficking to the distal dendrite of all cilia proteins (41). In *daf-19(m86)* mutants, GCY-22-wrmScarlet appeared enriched in an elongated ectopic membrane compartment protruding from the distal dendrite which was still in contact with AMsh but lacked any cilium (Figure 4A). The export of GCY-22-wrmScarlet to AMsh still occurred from this ectopic compartment and the number of GCY-22-wrmScarlet vesicles in AMsh was increased respective to wild type (Figure 4B), suggesting the cilia structure itself is not necessary for ectocytosis to occur. We next asked whether ectocytosis increased in dysfunctional cilium mutants. CHE-3 is a dynein heavy chain involved in cilia retrograde IFT. *che-3(cas511)* mutants display truncated cilia with bulbous endings (42–44). In *che-3(cas511)* mutants, GCY-22-wrmScarlet was enriched in a strongly misshaped ASER cilium (Figure 4A). The number of GCY-22-wrmScarlet carrying EVs in AMsh was increased, suggesting impaired retrograde IFT increases ectocytosis (Figure 4B). The BBSome complex subunit BBS-8 is dispensable for cilia assembly *per se* but is required for cilia function by regulating protein trafficking in and out of *C. elegans* cilia (45, 46). In addition, *bbs-8(nx77)* were shown to accumulate more EVs in the enlarged lumen of the male amphid sensilla (47). In *bbs-8(nx77)* mutants, ASER cilium length was more variable, the cilium base area was reduced (Figure 4A, C-D), and the number of GCY-22-wrmScarlet vesicles in AMsh was increased compared to controls (Figure 4B). This latter observation confirms that lack of BBS-8 activity promotes ectocytosis. Last, we asked if ectocytosis requires sensory activity to occur in cilia. To answer this question, we used mutants for TAX-4, a cyclic nucleotide-gated channel subunit expressed in ASER and required for its sensory functions (48). In *tax-4(p678)* mutants, the cilium length was slightly reduced, as previously observed for AWB fan-shaped ends (49) (Figure 4A and 4C). However, the number of GCY-22-wrmScarlet positive vesicles in the AMsh cell body was not modified in *tax-4* mutants (Figure 4B).

**Figure 4:**
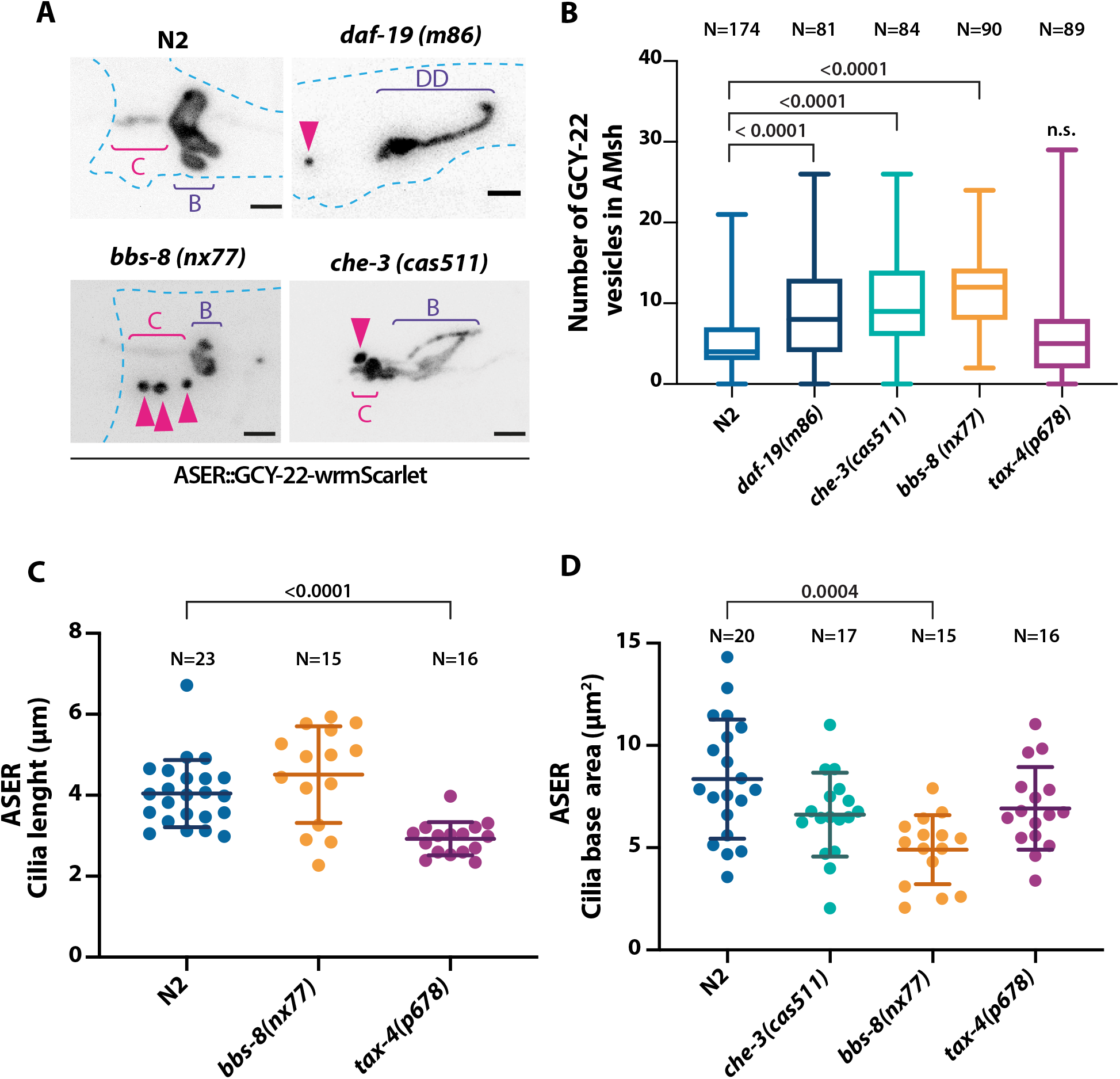
Ectocytosis of GCY-22-wrmScarlet is increased in bbs-8, che-3 and daf-19 mutants. A) Representative images depicting GCY-22-wrmScarlet distribution in ASER ciliary region for N2 and different mutants: daf-19, bbs-8 and che-3. In N2, GCY-22-wrmScarlet accumulates in the cilium (C) and cilia base (B). In daf-19 mutants, ASER cilium is missing. GCY-22-wrmScarlet accumulated in an elongated ectopic compartment arising from ASER distal dendrite (DD), an EV is observed in its vicinity within AMsh cytoplasm (magenta arrowhead). In bbs-8 mutants, cilia shape is not affected but we observed more EVs in AMsh cytoplasm in the vicinity of ASER cilium (magenta arrowheads). Cilia shape of che-3 is heavily disrupted, shortening ASER cilium and producing irregular cilium base shapes. Scale bar: 5 μm. B) Number of large ectosomes containing GCY-22-wrmScarlet within AMsh cell. Box and whiskers plot represents their median number, the interquartile range and the min/max values in N2, daf-19, bbs-8, che-3, tax-4. The number of vesicles is increased in daf-19, che-3 and bbs-8 mutants. Brown-Forsythe ANOVA, multiple comparisons corrected by Dunnett’s test. C) The ASER cilia length was evaluated based on GCY-22-wrmScarlet staining of the cilium. Cilia length is reduced in tax-4 and highly variable in bbs-8 mutants. Brown-Forsythe ANOVA, multiple comparisons corrected by Dunnett’s test. D) The ASER cilium base area is evaluated based on GCY-22-wrmScarlet staining in 2D projections. Cilia base area is reduced in bbs-8 mutants. Brown-Forsythe ANOVA, multiple comparisons corrected by Dunnett’s test. Scale bar: 5 μm.

Altogether these results suggest ectosome biogenesis does not require cilia, but likely stems from the strong directional traffic toward the cilia. The production and capture of ectosome carrying GCY-22-wrmScarlet is strongly increased in mutants regulating retrieval by the IFT*: bbs-8* and *che-3*, suggesting that lack cilia retrieval promotes ectocytosis. Finally, ectosome biogenesis was not modulated by sensory activity in ASER.

### Phagocytic activity in AMsh is required to maintain sensory cilia shape and function

The average size of the ectosomes carrying GCY-22-wrmScarlet (~1um) and their fast engulfment by AMsh suggests phagocytic mechanisms are involved. Using the export of GCY-22-wrmScarlet EVs from ASER to AMsh, we tested two genes involved in phagocytosis: *rac-1* and *dyn-1*. A hypomorphic mutation in *ced-10(n3246)* – the *C. elegans* ortholog of RAC1 – disturbs actin remodelling during the formation of phagocytic cup (50). Dynamin 2 is recruited early during phagosome formation and contributes to phagosome scission from plasma membrane (51). In human macrophages, the Dynamin 2 dominant negative mutation (K44A) abolishes GTP binding activity and disturb pseudopod extension (52). Similar dominant mutations in its *C. elegans* ortholog *dyn-1(G40E and K46A)* disturb engulfment rate, RAB-5 recruitment, phagosome maturation and degradation (53, 54). To cell-specifically assess the role of AMsh phagocytic activity, we expressed the dominant negative DYN-1(K46A) in AMsh only.

The ASER cilium shape was disturbed in *ced-10(n3246)* mutants as well as in animals expressing DYN-1(K46A) in AMsh (Figure 5A). In both manipulations, we observed abnormal tubulated protrusions attached to ASER cilia base, as if ectosome scission was slowed down. Nonetheless, *ced-10(n3246)* maintained normal cilium length and cilium base area compared to controls while AMsh::DYN-1(K46A) weakly reduced them (Figure 5A-C). The number of GCY-22-wrmScarlet positive EVs in the AMsh cell body was reduced in *ced-10(n3246)* mutants and was increased in animals expressing DYN-1(K46A) in AMsh (Figure 5D). Therefore, disrupting phagocytic activity in *ced-10(n3246)* mutants decreased EVs export to AMsh. Disrupting AMsh phagocytic activity cell-specifically by the expression of DYN-1(K46A) dominant negative increased ectosome uptake or reduced phagosome clearance cell-autonomously, and also disturbed ASER cilium shape non-cell autonomously. To confirm the non-cell autonomous effect AMsh phagocytic activity on ASER cilium morphology, we expressed DYN-1(K46A) in AMsh and the cytoplasmic mKate in ASER. Again, we observed abnormal tubulated protrusions still attached to the ASER cilia base that were not observed in controls (Figure 5E). The cilium length was reduced (Figure 5F) but the cilium base area size was not affected (Figure 5 – figure supplement 1 A-B). Many tiny vesicles were observed in the vicinity of ASER cilium, sometimes generating string of pearls within AMsh cytoplasm (Figure 5- figure supplement 1A’). To assess this non-cell autonomous effect of the expression of DYN-1(K46A) in AMsh on other amphid neurons, we examined the morphology of AFD NREs. In agreement with ASER results, we saw that NRE morphology of AFD was also affected: 25% animals displayed defects in AFD base morphology, 32% of the animals displayed enlargement of microvilli together with cilia base alterations, 35% showed a reduced number of microvilli and only an 8% showed a wild type phenotype (Figure 5G), overall demonstrating that disruption of AMsh phagocytosis has a detrimental effect on NRE shape.

**Figure 5.**
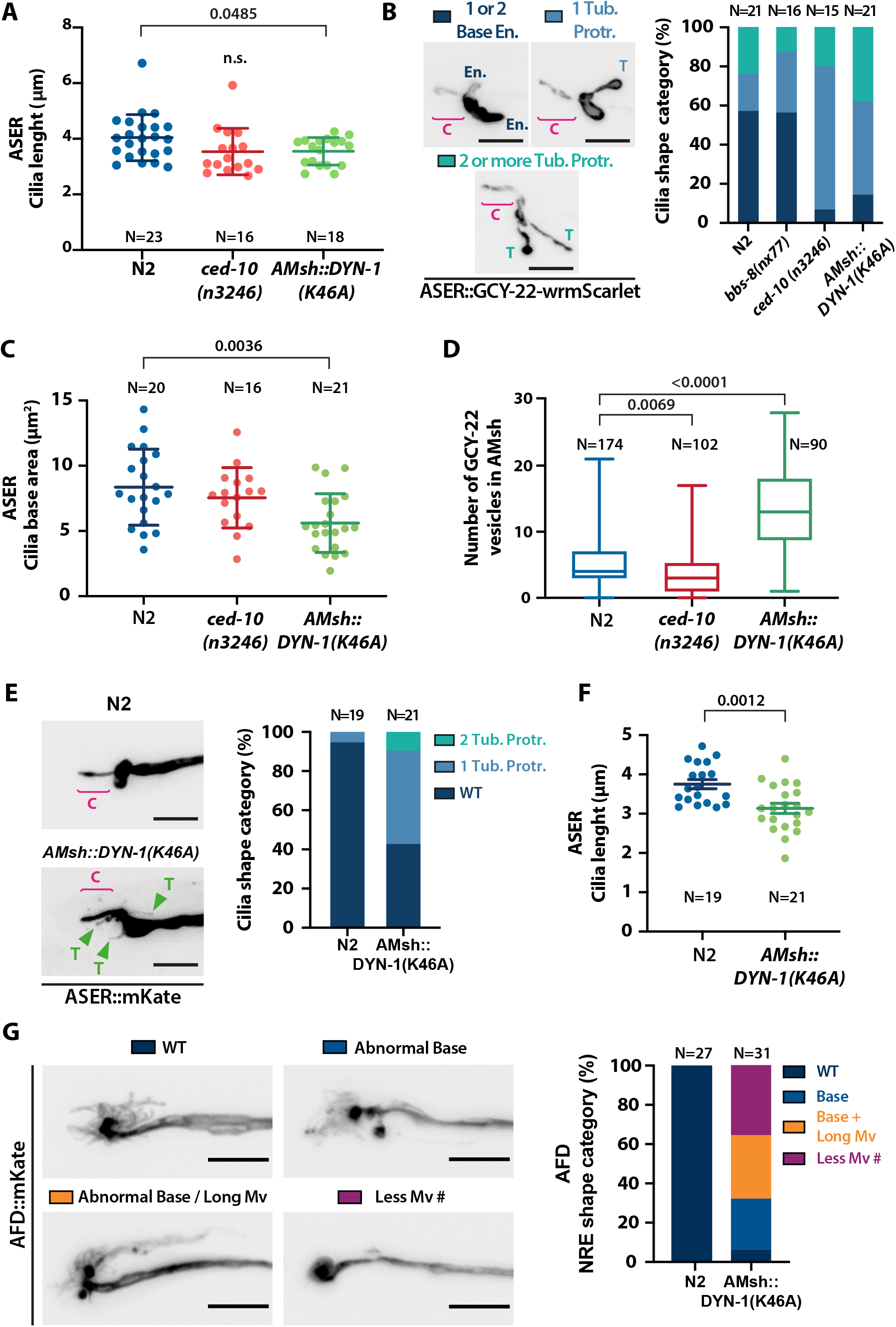
AMsh glia phagocytic activity is required to maintain a proper sensory cilia structure. A) The ASER cilia length was evaluated based on GCY-22-wrmScarlet staining of the cilium. Cilia length is slightly reduced in AMsh::DYN-1(K46A) transgenics and unaffected in ced-10(3246) mutants. Brown-Forsythe ANOVA, Multiple comparisons were corrected by Dunnett’s test. B) We examined ASER cilia shape in N2, ced-10(n3246) mutants and AMsh::DYN-1(K46A) transgenics animals expressing GCY-22-wrmScarlet in ASER. Three categories were made according to ASER cilia shape: animals displaying 1 or 2 cilia base enlargements still connected to PCMC without signs of tubulation, animals displaying 1 tubulated protrusion or animals displaying 2 or more tubulated protrusions. (C: cilium, En.: base enlargement, T: tubulated protrusion). The percentage of each cilium shape category is given for each genotype. Ced-10(n3246) and AMsh::DYN-1(K46A) transgenics strongly increase the number of animals showing tubulated protrusions. C)The cilia base area is evaluated based on 2D projections. The PCMC area is not significantly modified in ced-10 mutants but reduced in AMsh::DYN-1(K46A) transgenics. D) Number of large ectosomes containing GCY-22-wrmScarlet within AMsh cell for N2, ced-10(n3246), AMsh::DYN-1(K46A). Ectosome number in AMsh is decreased in ced-10(n3246) mutants and increased AMsh::DYN-1(K46A) transgenics. Brown-Forsythe ANOVA, multiple comparisons corrected by Dunnett’s test. E) We examined ASER cilium shape in N2, ced-10(n3246) mutants and AMsh::DYN-1(K46A) transgenics animals expressing mKate. The categories for ASER cilium shape are: wild type phenotype (WT), animals displaying 1 tubulated protrusion or animals displaying 2 or more tubulated protrusions. (C: cilium, T: tubulated protrusion). AMsh::DYN-1(K46A) transgenics strongly increase the number of animals showing tubulations (green arrowheads). F) The ASER cilia length was evaluated based on mKate staining of the cilium. Cilia length is slightly reduced in DYN-1(K46A) transgenics. Brown-Forsythe ANOVA, Multiple comparisons were corrected by Dunnett’s test. G) NRE shape of AFD in N2 and in AMsh::DYN-1(K46A) transgenics expressing mKate in AFD. 4 categories were established: wild type phenotype (WT), abnormal AFD cilia base (Base), abnormal base + elongated microvilli (Base + Long Mv) and reduced number of microvilli (Less Mv #). Scale bar: 5 μm.

We reasoned that the aforementioned abnormal cilia structure in AMsh::DYN-1(K46A)-expressing animals could lead to defects in the sensory perception of the amphid sensory neurons. We explored this possibility by using two well-described chemotactic assays for the chemosensory neurons ASER and ASH (55). *C. elegans* shows a preference for NaCl concentrations associated to their cultivation on food, a behaviour that depends on ASE neurons. Within a linear NaCl, gradient we observed a loss of attraction for the NaCl concentrations associated to food in animals expressing DYN-1(K46A) in AMsh (Figure 6A and Figure 6 – figure supplement 1A). When placed into a gradient of the aversive Copper ions (Cu^2+^), *C. elegans* navigate towards low concentrations, a behaviour that depends on the ASH nociceptor neurons. Within a linear Cu^2+^ gradient, animals expressing DYN-1(K46A) in AMsh completely lost repulsion to Cu^2+^ (Figure 6B and Figure 6 – figure supplement 1B). These results imply that ASH and ASER sensory function is altered non-cell autonomously by expression of DYN-1(K46A) in AMsh. To confirm that ASH was otherwise functional, we used ChannelRhodopsin (ChR2) to stimulate ASH independently of any sensory cues and sensory machinery. We did not observe any abnormal response to blue light exposure in animals expressing DYN-1(K46A) in AMsh (Fig 6C). Therefore, disturbing glial phagocytic function affects cilia shape and sensory perception, but downstream signal transduction, including sensory neuron depolarisation, neurotransmission and signal integration by the nervous system remains unaltered.

**Figure 6.**
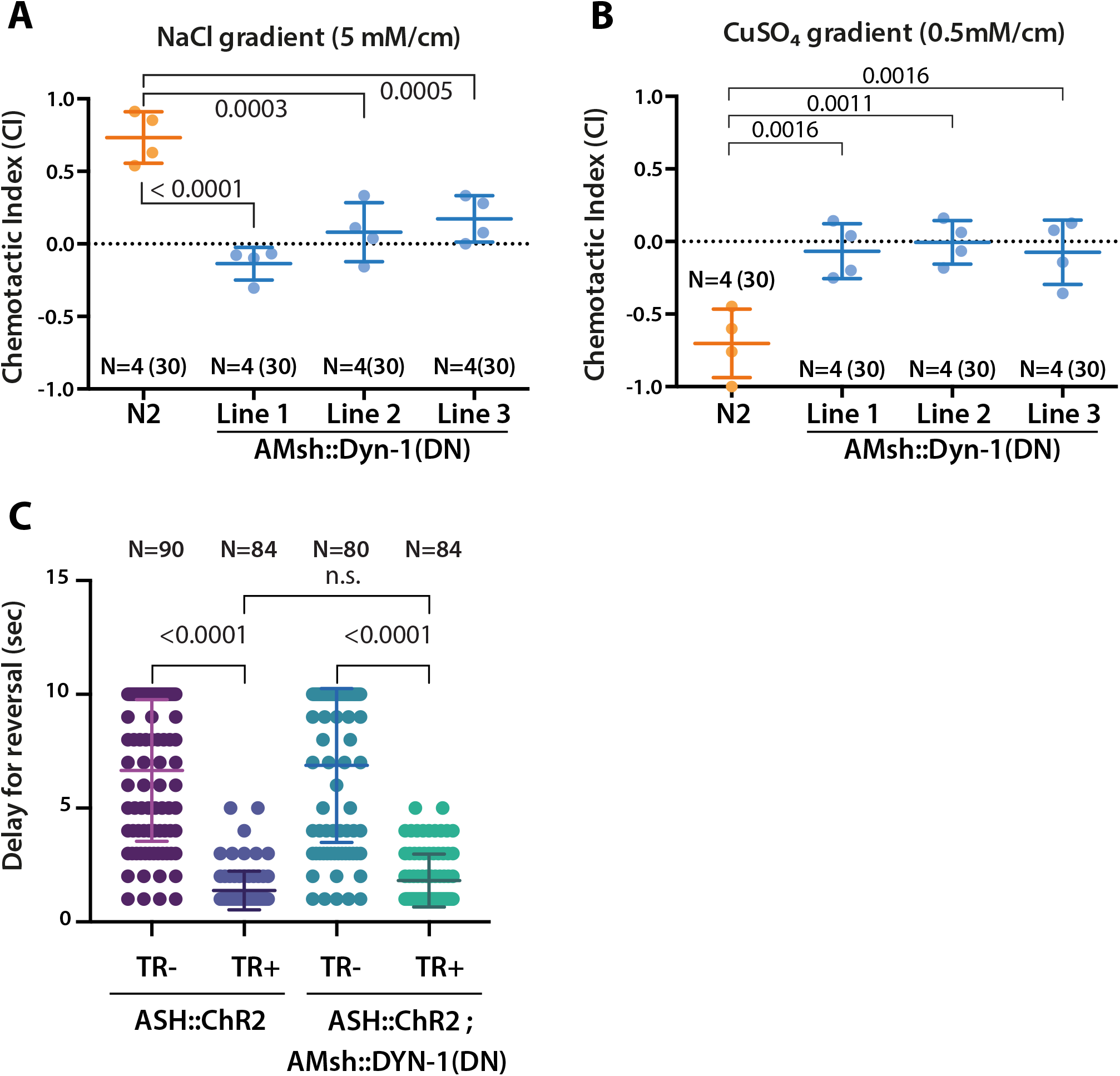
AMsh phagocytic activity is required for sensory functions. A) Chemotactic indexes of 4 independent assays made with 30 N2 or AMsh::DYN-1(K46A) transgenics in linear 5 mM/cm NaCl gradients. One-way ANOVA, multiple comparisons corrected by Tukey test. B) Chemotactic indexes of independent 4 assays made with 30 N2 or AMsh::DYN-1(K46A) transgenics in linear 5 mM/cm CuSO_4_ gradients. One-way ANOVA, multiple comparisons corrected by Tukey test. C) Transgenic animals expressing ChR2(H134R) in ASH sensory neurons (ASH::ChR2(H134R); lite-1) exhibit fast reversal (minimum 1~2 backward head swings) in response to blue light exposure (15mw/mm^2^). Expression of AMsh::DYN-1(K46A) does not modify this avoidance response. Kruskal–Wallis test, multiple comparison corrected by Dunn’s test.

### Pruning by AMsh glia is not necessary for the production of EVs by AFD

In several animal models, glia plays an active role in the fine sculpting of the nervous system. Glial cells can prune excess synapses, axonal projections, dendritic spines and sensory endings of neurons (56). Pruning of the glia-embedded receptive ends of AFD, AWA, AWB AWC by AMsh was suggested before: ablation of AMsh as well as mutants affecting AMsh properties alter their NRE structure (57, 58). Similarly, AMsh expression of DYN-1(K46A) modified AFD receptive endings. Therefore, we explored a potential role for pruning to explain the export of TSP-7-wrmScarlet from AFD to EVs in AMsh. First, time-lapse recordings from animals expressing TSP-6-wrmScarlet in AFD neurons showed many of the TSP-6-wrmScarlet EVs seem to originate from AFD microvilli endings. Additionally, we could observe elongation and retraction of microvilli (Video 7). We first asked whether presence of AFD microvilli was strictly necessary to observe export of TSP-7-wrmScarlet from AFD to EVs in AMsh. To this end, we used a mutant for TTX-1, a transcription factor required for correct morphogenesis of AFD terminals. *ttx-1(p767)* mutants are defective for AFD microvilli formation but maintain their cilia (5, 59). In *ttx-1(p767)* mutants, TSP-7-wrmScarlet remained enriched in AFD receptive endings lacking microvilli (Figure 7A). TSP-7-wrmScarlet was still exported to AMsh in the *ttx-1(p767)* although the export was strongly reduced (Figure 7A-B). Therefore, the AFD microvilli are important but not absolutely required for TSP-7-wrmScarlet export from AFD to EVs in AMsh. We next asked whether the export of TSP-7-wrmScarlet from AFD to EVs in AMsh required the full amphid sensilla structure or only the close apposition of AFD to AMsh. To answer this question, we used the *dyf-7(m537)* mutants where the AMsh still ensheathes AFD nerve receptive endings but in an ectopic location inside the head (60). Despite this displacement of AFD nerve receptive endings in *dyf-7* mutants, the transfer of TSP-7-wrmScarlet from AFD to EVs in AMsh still occurred: we observed the typical fluorescent EVs throughout the cell body of AMsh (Figure 7C). Therefore, export of TSP-7-wrmScarlet only requires the AFD receptive endings to contact AMsh, independently of a proper morphology and location of the amphid sensilla. We next asked whether the presence of AMsh is required for the production of TSP-7-wrmScarlet EVs by AFD. For this, AMsh was genetically ablated by means of Diphtheria Toxin (DTA) expression. In the strain we used (OS2248), amphid neurons and their receptive endings properly develop before AMsh ablation occurs in late embryogenesis (57). In absence of AMsh, TSP-7-wrmScarlet export still occurred from AFD receptive endings to EVs located in other neighbouring cells, presumably hypoderm cells from the head (Figure 7D, right panels). Therefore, pruning of AFD microvilli by AMsh or an instructive signal from AMsh is not strictly necessary for the production of TSP-7-wrmScarlet EVs by AFD. Instead, the export of TSP-7-wrmScarlet simply requires the EVs produced by AFD to be captured by any competent neighbouring cell type. Among the local cells undergoing constitutive endocytic activity and located in the proximity of AFD NREs are all the ciliated amphid neurons. Interestingly, in absence of AMsh, a fraction of TSP-7-wrmScarlet is exported from AFD to EVs located in the cytoplasm of a subset of ciliated amphid neurons also expressing the osm-3p::mEGFP transgene (Figure 7 – figure supplement 1E-H). Therefore, amphid neurons can also take up TSP-7-wrmScarlet at their receptive endings in absence of AMsh.

**Figure 7:**
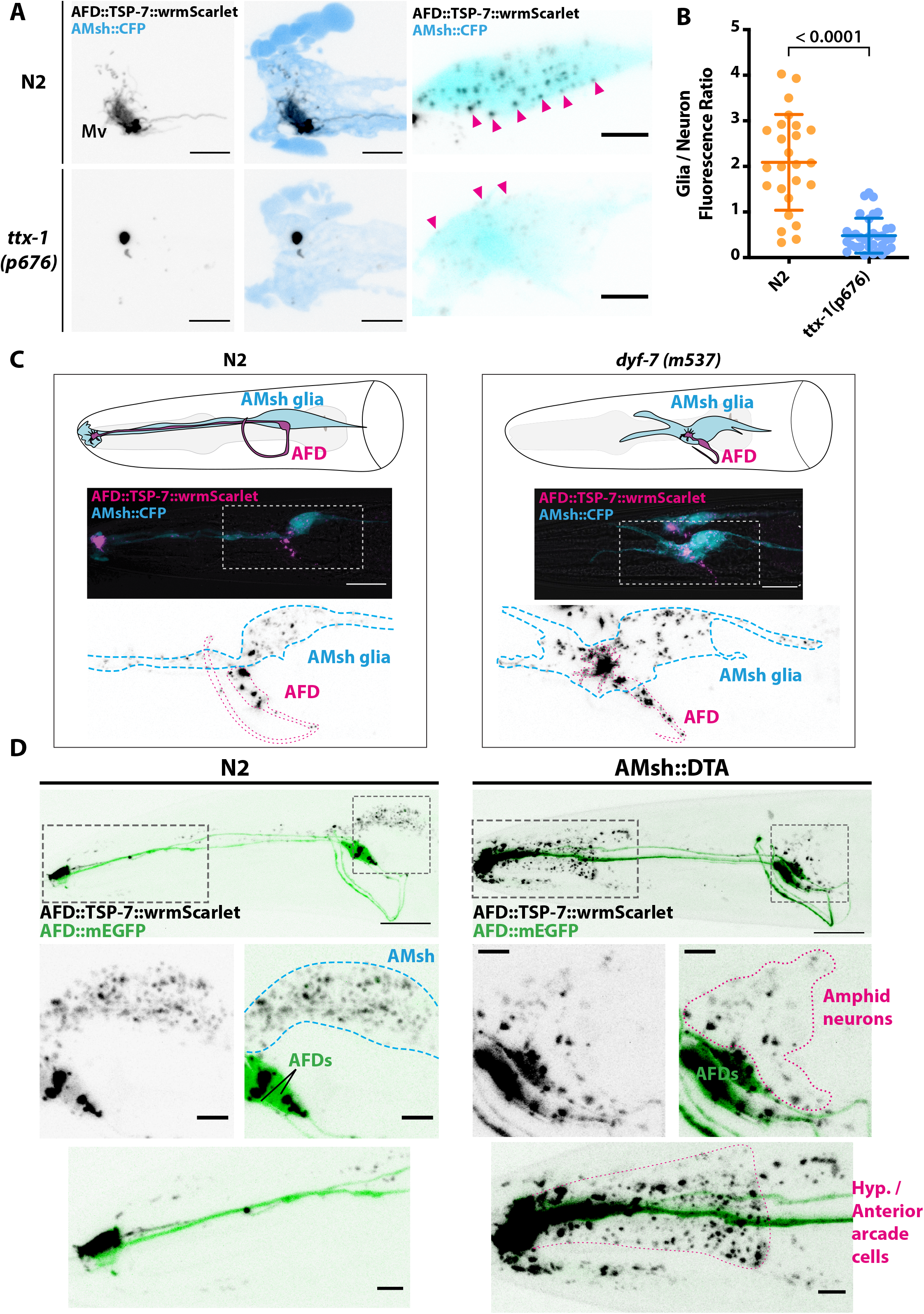
Position and presence of glia are not necessary for EV production and export to occur. A) In ttx-1(p767) mutants, the microvilli (Mv) disappear but TSP-7-wrmScarlet remained enriched in the remaining AFD receptive end. Scale bar: 5 μm. B) TSP-7-wrmScarlet export was quantified by measuring AMsh fluorescence intensity and normalised to the AFD cell body intensity. TSP-7-wrmScarlet export is decreased in ttx-1(p767) mutants. Unpaired t test with Welch’s correction. C) In dyf-7(m537) mutants, TSP-7-wrmScarlet remained enriched in AFD receptive endings although the receptive ending was displaced posteriorly in the animal’s head but still embedded within AMsh. TSP-7-wrmScarlet was still exported to AMsh cell body in a similar manner to wild type controls. Scale bar: 20 μm. D) Representative images displaying differential tissue capture of EVs when glia is ablated genetically post-embryogenesis. Animals expressed AFD::mEGFP and AFD::TSP-7-wrmScarlet. TSP-7-wrmScarlet is enriched in AFD receptive end in both experimental conditions. EVs are exported to AMsh in control conditions. In absence of glia, EVs containingTSP-7-wrmScarlet were still produced but were exported to large cells at the surface of the nose, likely the hypodermal cells. TSP-7-wrmScarlet was also exported to amphid sensory neurons. Scale bar: 20 μm for top head images, 5 μm for insets.

Our observations suggest EVs produced by AFD are physiologically taken up by the scavenging activity of AMsh as long as AFD receptive ends are embedded within AMsh. The phagocytic activity of AMsh contributes to define the size and shape of AFD (and other amphid neurons) receptive endings. However, the absence of AMsh pruning activity or AMsh-derived signal does not suppress EV production, suggesting an intrinsic production mechanism of EVs by AFD. In absence of AMsh, any local endocytic activity can take up these EVs.

## Discussion

Several early arguments suggested that EVs produced by *C. elegans* males correspond to ectosomes shed from the cilia of the male neurons (17). While writing, new observations suggested that ciliary ectosomes may be shed from two sites of the cilia of CEM neurons (61). Our results confirm and extend these conclusions: many - potentially all- hermaphrodite ciliated neurons produce ciliary ectosomes from their cilia. Using in vivo recordings and volumetric confocal acquisitions, we could identify the location for the biogenesis and excision of ciliary ectosomes. The fate of these ectosomes varies according to the location of their biogenesis: ectosomes budding from the cilia base are simultaneously captured by the contacting glial cells, while ectosomes shed from the cilia tip were environmentally released, exiting through the pore of the sensilla (Figure 8). Ectosomes budding from ciliary tip subcompartment were observed for the ASER, ASH/ASI, and IL2 neurons. Ectosomes budding from the ciliary base and simultaneously captured by their contacting glial cells were observed across all the neurons analysed in this study, indicating that most ciliated neurons are competent to form ectosomes from their cilia base. Within these glial cells, the ectosomes are directed to acidic organelles that likely belong to the endolysosomal pathway (Figure 8). All cilia protein assayed in this study were fused to wrmScarlet: an RFP-derived fluorophore noticeably more tolerant to acidic pHs than its GFP-derived counterparts (62, Shinoda, 2018 #85). This observation, or the larger lumen of the male cephalic sensilla might explain why export of GFP-tagged cilia proteins to the supporting glia was not observed previously. While the export of ciliary cargoes to the glia suggests ciliary EVs represent a form of cellular disposal, several examples of bioactive ciliary EVs carrying information across cells are reported in *C. elegans* and other species (19). It is therefore possible that ciliary ectosomes exported from sensory neurons to AMsh has additional significance in cell-to-cell communication between neuron and glia. For example, *C. elegans* mutants defective in ciliogenesis also show a defect in AMsh membrane trafficking leading to enlarged amphid pore lumen (5). This correlation suggests an interesting possibility where neuronal ectosomes could signal the cilia state to AMsh, while the latter responds by secreting factors in the vicinity of neuronal receptive endings (47, 49, 63).

**Figure 8.**
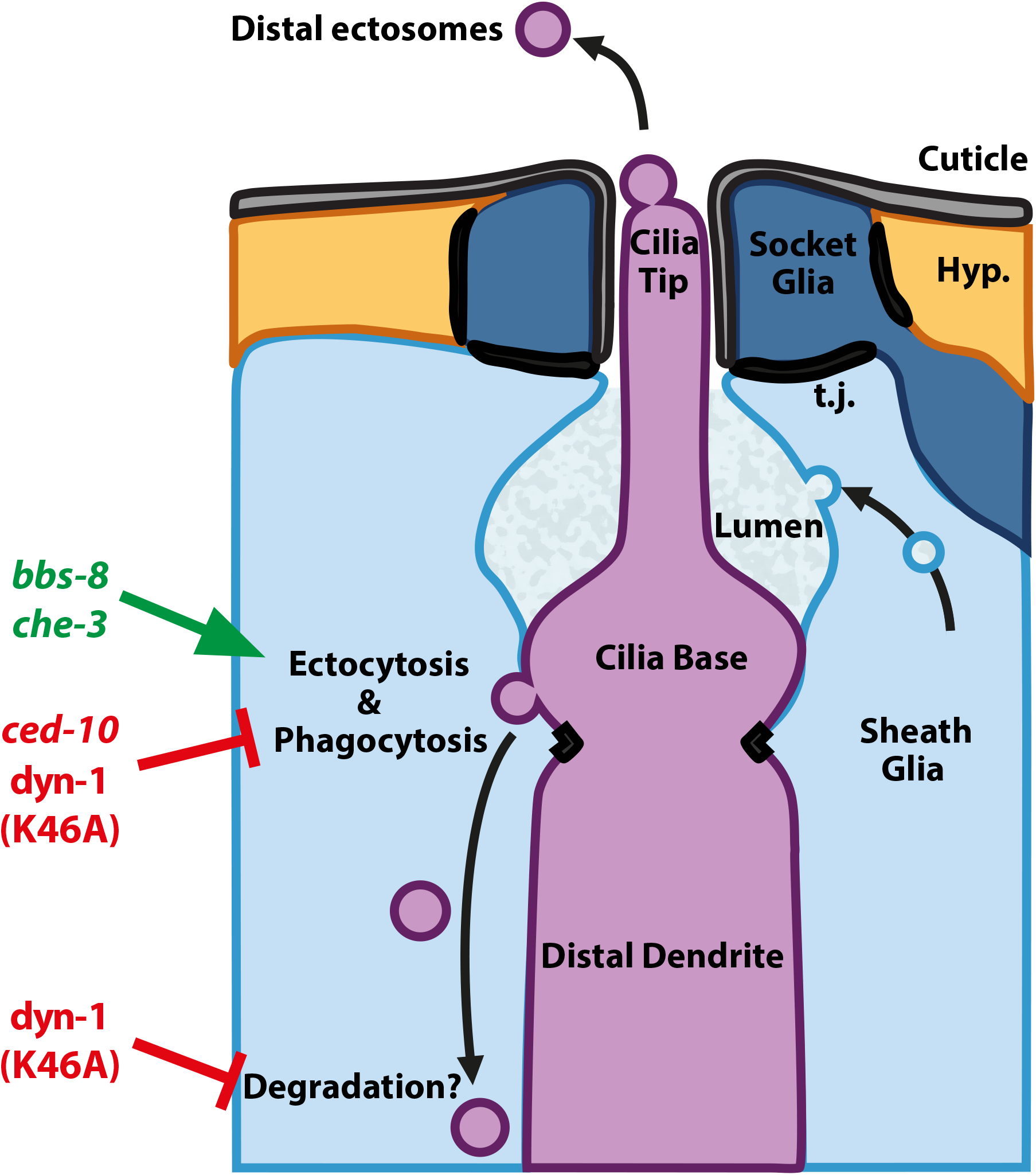
Mechanisms underlying ectocytosis from ciliated neurons. We suggest a model where ectocytosis is inherent to cilia and where neurons and glia cooperate to readily remove basal ectosome from ciliary membranes (magenta). Ectocytosis occurs from 2 different ciliary locations: the cilia tip and the cilia base. When ectosomes are shed from the cilia tip they are environmentally released. Sheath glia (light blue) embeds the ciliary base of ciliated neurons, when ectosomes bud from the base they are concomitantly phagocytosed by their associated sheath glia. Besides its crucial role in secretion of extracellular matrix content, we suggest glia also plays an important function to maintain cilia structure and composition and to recycle ectocytosed material. Mutations in genes involved in cilia protein retrieval by IFT, like bbs-8 and che-3, fail to balance import and removal of ciliary proteins and show increased export and ectocytosis events to the sheath glia. A ced-10 mutation reducing phagocytic activity decreases the uptake of ectosomes and modifies cilia shape. Cell-specific manipulations of AMsh glia phagocytic activity by expression of dyn-1 dominant negative transgene suggest that a tight regulation of sheath glia phagocytosis contributes to shape the nerve receptive endings. In addition, dyn-1 manipulation also reduces phagosome degradation.

What is the purpose of ciliary ectocytosis? Cilia size, composition and signalling are tightly regulated (64). For example, tuning of the Hedgehog signalling is achieved by controlling cilia entry, removal and disposal of Hedgehog effectors (65–67). The best-known mechanisms for entry and removal of proteins from cilia are mediated by the IFT cargo adapters IFT-A and BBSome, respectively. More recently, other mechanisms than BBSome retrieval were described to remove proteins from cilia, including endocytosis at periciliary membrane and ciliary ectocytosis (18, 68). In IMCD3 kidney cell line, ciliary ectocytosis was shown to provide an additional route to remove signalling receptors from cilia (18). In these cells, cilia defective for BBSome retrieval accumulated receptors in their cilia and increased their removal by ectocytosis (18, 69). Therefore, ciliary ectocytosis may provide a safeguard measure to maintain an appropriate cilia composition in absence of receptor retrieval by IFT. Our results are consistent with this hypothesis: in *bbs-8* and in *che-3* mutants where IFT retrieval of GCY-22-wrmScarlet is reduced, we observed an increased release of GCY-22-wrmScarlet-containing ectosomes (39, 70) (Figure 8). Previously, *bbs-8* mutants were shown to accumulate EVs in an enlarged lumen of the amphid sensilla (47). However, the authors considered these EVs observed in *bbs-8* as pathological, since they were not observed in the amphid lumen of control animals. Instead, our results suggest ectosomes are physiologically produced from amphid neurons in controls but are directly captured by AMsh and therefore not observed in the lumen of control amphids. Surprisingly, we observed that cilia themselves are not strictly necessary for ectocytosis: *daf-19* mutants, which are void of any ciliary structure, still exported GCY-22-wrmScarlet to EVs in AMsh. It is therefore likely that ectocytosis is a by-product of the strong and polarised traffic of GCY-22-wrmScarlet towards the distal dendrite. As previously suggested, we hypothesise that ectocytosis represents one mechanism to balance out the continuous import of membrane and membrane proteins to cilia (64).

Does ciliary ectocytosis occur in physiological conditions? Physiological production of ciliary ectosome by neurons is well documented in *C. elegans* male ciliated neurons as well as in human photoreceptor (17, 71). We demonstrate that lipophilic DiI is captured by ciliated amphid neurons and exported from them to AMsh by ciliary EVs. Importantly, this assay shows that export of ciliary membrane occurs in absence of protein overexpression, arguing for a physiological production of ciliary EVs. Also, cell-specific manipulation of glial phagocytosis induced abnormal cilia structure, suggesting that physiological production of EVs is cleared by the glial cells. However, we do not exclude that overexpression of ciliary membrane proteins could stimulate production of ectosomes by neurons and/or their capture by glial cells. One possibility would be a saturation of the IFT retrieval machinery by overexpression of ciliary membrane proteins. Cargo accumulation might also play an active role in ectosome formation either by instructing a sorting machinery or by changing the membrane properties. For example, clustering of membrane proteins with intrinsic conical shape creates microdomains which can bend their associated membranes (72). Tetraspanins were previously suggested to contribute to EVs biogenesis and cargo sorting to the EVs (73). TSP-6 overexpression might therefore facilitate EV biogenesis by its effect on protein clustering, membrane curvature and/or its sorting to high curvature regions of the plasma membrane as suggested for its mammalian ortholog, the tetraspanin CD9 (74).

How does cargo enter ectosomes? Although ciliary ectosomes derive from cilia membrane, their protein composition is different (75). In IMCD3 cilia, activated GPCRs are selectively sorted to ciliary EVs, leaving most of the non-activated GPCRs behind (18). In contradiction with these results suggesting a highly selective sorting machinery, all the ciliary protein we assessed -except SRBC-64- were able to enter basal ectosomes. In addition to a sorting machinery, cargo themselves can have an active role in EV biogenesis by their bending properties, their interactions with bending proteins and/or their preferential sorting to bending membranes. In agreement to this hypothesis, we show that EV composition correlates with changes in EV diameter. The presence of GCY-22 increased the diameter of EVs carrying TSP-6, while presence of TSP-6 reduced the diameter of the EVs carrying GCY-22 (Figure 3 – Supplement 1B-D). This observation suggests cargo interact within the plasma membrane to define the EVs size. For example, GCY-22 overexpression might neutralise the bending effects of TSP-6, leading to larger EVs. Once membranes start bending, curvature-dependent sorting of cargo can contribute to their trafficking, as recently observed for sorting of activated GPCRs (76). We also show that sorting to basal or distal ectosomes rely on cilia trafficking biases causing local enrichment/accumulation of cargoes in ciliary sub-compartment, e.g. TSP-6/TSP-7-wrmScarlet is enriched in IL2 cilia base, and mostly released in basal ectosomes. However, we observed that TSP-6 and TSP-7-wrmScarlet were also able to enter distal ectosomes, but only when they were present at the IL2 cilia tip during distal ectosome formation (Video 4). Therefore, cargo entry in basal and/or distal ectosomes does not result from a selective sorting but rather results from their basal and/or apical enrichment when ectosome budding occurs. Sensory cilia of *C. elegans* vary in term of axonemal structure, microtubule arrangements, post-translational modifications of tubulin as well as IFT motors composition (5, 22, 77, 78). These cilia specificities contribute to determine whether cargoes are enriched in distal or basal cilia in a given cell type and can explain the high propensity of male ciliated neurons to produce distal ectosomes carrying PKD-2.

Does ectocytosis maintain cilia shape and function? A preprint of Raiders et al. showed that CED-10 activity in AMsh dictates AFD microvilli engulfment rate similarly to what we observed for DYN-1(K46A) (58). However, we show that EVs remained produced by AFD receptive endings in absence of AMsh glia; these EVS were alternatively taken up by hypodermal and neuronal cells. Taken together, these results suggest a model where glial phagocytic activity is secondary to the inherent formation of EVs by NREs. Because the sheath glia embeds partially or totally the NREs, glial phagocytosis often goes hand in hand with ectocytosis events. Neurons and glia would therefore cooperate to readily remove and recycle ciliary membranes (Figure 8). Besides its crucial role in secretion of extracellular matrix to the sensory pores, we suggest sheath glia also plays an important function to maintain cilia structure and composition and to recycle material discarded by its associated neurons. When AMsh phagocytic activity is disturbed in *ced-10* mutants or in glia-specific *dyn-1* dominant negative transgenics, the morphology of ASER and AFD receptive endings was severely altered. Not surprisingly, disrupted cilia morphology correlates with abnormal ASER and ASH sensory responses. Therefore, removal of receptors via ectocytosis and their capture by the sheath glia might contribute to the sensory function of the *C. elegans* cilia. Similarly, the mammalian photoreceptor neurons coexist with closely apposed Retinal Pigmented Epithelium (RPE), which phagocytoses a packet of ~100 discs shed daily from the cilia tip of each photoreceptor (2). Mutations in MERKT, a transmembrane protein involved in recognition and phagocytosis of the photoreceptor outer segments by RPE, leads to accumulation of debris and result in loss of vision and photoreceptor degeneration (79).

We show that ectocytosis is constitutive of *C. elegans* sensory cilia and contributes to grant appropriate structure and function. *C. elegans* ciliary ectosomes were previously shown to facilitate interindividual communication (17). The export of ciliary ectosome from neurons to glia suggest these might also contribute to cell-cell communication. Ectosome release was shown to involve ciliary accumulation of PI(4,5)P2 and subsequent actin polymerisation in cilia (18, 80). However, the cargo sorting machinery remains largely unknown. Here we establish a model to address these questions.

## Materials and Methods

### Strain and genetics

*C. elegans* were cultured on NGM agar plates provided with E.coli OP50 bacteria and grown under standard conditions unless otherwise indicated (81, 82). All strains were grown at 20°C unless otherwise indicated. Strains used in this study are listed in Supplementary Table 1.

### Molecular biology and transgenic strains

*C. elegans* N2 genomic DNA was used as template for cloning PCRs. Cloning PCRs were performed using Phusion High Fidelity DNA polymerase (M0530L, New England BioLabs) and then validated by Sanger sequencing. Cloning information is provided in Supplementary List 2. *C. elegans*-optimized wrmScarlet was a gift from Thomas Boulin (62).

Plasmids used to generate transgenic animals were generated by means of Multisite Three-Fragment Gateway® Cloning (Invitrogen™, Thermo Fisher Scientific, MA, USA). In brief, PCR fragments containing AttB recombination sequences were recombined into DONOR vectors by means of BP Clonase reactions (Invitrogen™, Thermo Fisher Scientific, MA, USA), effectively creating a collection of ENTRY clones. Three entry clones, (Pos1 + Pos2 + Pos3), and a destination vector, were used to create a pEXPRESSION vector. The destination vector used in the constructs for this study was modified from the pDEST™ R4-R3 Vector, a 3’UTR sequence of the *let-858* gene was added between attR3 and AmpR. Plasmids used in this study are available in Supplementary Table 3.

Site-specific mutagenesis for DYN-1(K46A) dominant negative mutation was performed by standard PCR method using a plasmid with the wild type dyn-1 genomic sequence as template. Overlapping primers of ≈ 40bp were designed and PCR reaction was set for 25 cycles using Pfu Turbo polymerase. Following the reaction, mixes were digested with DpnI to remove bacterial methylated DNA and transformed into DH5-alpha competent bacteria. Plasmids DNA was extracted, and mutations were confirmed by sequencing of plasmid DNA at Eurofins Genomics (Ebersberg, Germany).

All transgenic worms were generated by microinjection with standard techniques (83). For most injected constructs, injection mixes were composed of 30 ng/μl targeting constructs, 30 ng/μl of co-injection markers and 40 ng/μl of 1kb Plus DNA mass ladder (Invitrogen™, Thermo Fisher Scientific, MA, USA) as carrier DNA to have a final injection mix concentration of 100 ng/μl. For *tsp-6* constructs, a final concentration of 5 ng/μl of targeting construct was used. For mKate constructs, a final concentration of 15 ng/μl. Each promoter used was first tested for expression pattern in order to make sure it was only expressed in neurons and never in glial cells. Due to mosaicism generated by injection of unstable extrachromosomal arrays, 3 independent transgenic lines were generated for each of the constructs used in this study, only one strain was shown to be representative of the phenotype observed in the 3 independent transgenic lines.

### DiI staining

Animals were synchronized by selecting L4 larva the day prior to the assay. Worms were stained using a modified version of Wormatlas Anatomical Methods (84). We used the lipophilic DiI (1,1’-Dioctadecyl-3,3,3’,3’-Tetramethylindo carbocyanine Percholrate) (#42364, Sigma-Aldrich) to stain a subset of amphid sensory neurons (ASK, ADL, ASI, AWB, ASH and ASJ). Briefly, worms were washed from plates with M9 and placed on a 1.5 mL microcentrifuge tubes. Worms were washed twice with M9 afterwards by spinning them down at 3000 rpms to bring the worms to the bottom of the tube. Worms were treated with M9 or with M9 supplemented with 25mM NaN_3_ 15 min prior to staining (See Figure 1 - Figure Supplement 1A). 15 min of 25mM NaN_3_ were sufficient to fully anesthetize the animals as evaluated by the cease of pharyngeal pumping. DiI (2 mg/mL) stock solution was diluted 1:200 for staining. For staining, the worms were incubated for 20 minutes in the dark on a rocker at 30rpms in M9 or in M9 + 25mM NaN_3_. After staining, worms were either washed 3 times with M9 + 25mM NaN_3_ and directly mounted on a microscope slide for imaging or were left on OP50-seeded plates and allowed to recover from anesthesia for 1 hour. After 1 hour the recovered animals were directly mounted for imaging (see below).

### Image acquisition

Worms were synchronized either by bleaching a population of gravid worms or by an egg-laying window. Worms were reared at 20°C up to the right stage. All animals were imaged at Day 1 adulthood unless otherwise stated. Synchronized animals were mounted on 2% agarose pads and anesthetized with 25 mM Sodium Azide (NaN_3_)(#S2002,Sigma-Aldrich, MO, USA) dissolved in M9 solution. Images were acquired in the following 10-60 minutes after animals were anesthetized. Images were acquired at the Light Microscopy Facility LiMiF (http://limif.ulb.ac.be) at the Université Libre de Bruxelles, Faculté de Médecine, Campus Erasme, on a LSM780NLO confocal system fitted on an Observer Z1 inverted microscope. (Carl Zeiss, Oberkochen, Germany). Images in which the full animal’s head is displayed were acquired using a LD C-Apochromat 40x/1.1 W Korr M27 objective. The settings for these images were as follows: frame size was set at 1024 × 1024 pixels with a pixel size of 0.13 μm × 0.13 μm, pinhole size was set to 1 Airy Unit, Z-step optical sections varied across images (from 0.3-0.64 μm/step size) depending on the desired ROI volume (ranging from 10 to 35 μm in the Z axis), pixel dwell was set to 0.79 μs and averaging was set to 4. High resolution details (inset images) were acquired using an alpha Plan Apochromat 63x/1.46 Oil Korr M27 objective. The settings for these images were as follows: frame size was set to 412 × 412 pixels with a pixel size of 0.08 μm × 0.08 μm, pinhole size was set to 1 Airy Unit, Z-step optical sections varied across images (from 0.2-0.3 μm/step size) depending on the desired ROI volume (ranging from 10 to 20 μm in the Z axis), pixel dwell was set to 1.95 μs and averaging was set to 4. Images with a single channel were acquired using GaAsP detector. Images with multiple channels were acquired simultaneously using PMT detector for the following fluorophores: CFP or mEGFP, and the GaAsP detector for wrmScarlet and mCherry signals. For individual channel acquisitions, the Main Beam Splitter matched the excitation wavelength of each used fluorophore. For simultaneous CFP/wrmScarlet or mKate images a 458/543 Main Beam Splitter was used. For simultaneous mEGFP/wrmScarlet or mCherry a 488/543 Main Beam Splitter was used. The following fluorophores excitation (Ex) and detection wavelengths (DW) were used: for CFP (Ex: 458 nm – DW: 463-558 nm), for mEGPF (Ex: 488 nm – DW: 493-569 nm), for wrmScarlet/ mKate / DiI (Ex: 543 nm – DW: 570-695 nm), for mCherry (Ex: 543 nm – DW: 588-695 nm). Fluorophore excitation and detection wavelength ranges were set according to the information available FPbase database for each fluorophore (85). Laser power and detector gain settings were adjusted to maximize signal-to-noise ratio and minimize saturation when possible. Images were saved in .czi Zeiss file format.

Images used to quantify the number of GCY-22 ectosomes within AMsh were done using an Axioimager Z1 microscope. Images of the full animal’s head were acquired using a Plan Apochromat 20x/0.8 M27 objective and an AxioCam MR R3 camera (Carl Zeiss, Oberkochen, Germany). The settings for these images were as follows: frame size was set at 1388 × 1040 pixels with a pixel size of 0.323 μm × 0.323 μm. For CFP/mEGFP channel, the laser excitation was done using a 488 nm laser and using the following filters for excitation and wavelength detection (Excitation filter: 450-490 nm / Detection filter: 500-550 nm), exposure time was set to 20 ms. For wrmScarlet channel, the laser excitation was done using a 453 nm laser and using the following filters for excitation and wavelength detection (Excitation filter: 538-562 nm / Detection filter: 570-640 nm), exposure time was set to 600 ms.

### In vivo time-lapse imaging

Animals were synchronized as mentioned before and imaged at Day 1 adulthood. For time lapse imaging, a drop of anesthetic solution: 10mM Tetramisole Hydrochloride (#L9756, Sigma-Aldrich, St. Louis, MO, USA) in M9 was placed in a FluoroDish (FD35-100, World Precision Instrument, Inc., FL, USA). Several animals were placed in the drop for 10-15 min prior to imaging until anesthetized. Immobilized worms were then covered with a layer of 4% agarose and maintained in anesthetic solution throughout the acquisition duration. We used the LSM780NLO confocal microscope with either the LD C-Apochromat 40x/1.1 W Korr M27 objective or the alpha Plan Apochromat 63x/1.46 Oil Korr M27 objective to acquire the time-series images. Time intervals between frames were set as follows for each video: Video 1 (1.23 s), Video 2, 3 and 4 (940 ms), Video 5 (2.84 s), Video 6 (1.27 s), Video 7 (2.84 s)

A single focal plane representative of the NRE area was used to acquire the time-series. For Video 6 and 7, pinhole size was set to 2 AU to maximize signal. Acquisition settings were set as described previously.

### Image processing

Confocal images were processed using FIJI (86) and Zen 2.6 Pro (Blue edition) software (Carl Zeiss, Oberkochen, Germany). Z-stack acquisitions were converted into a 2D image using Maximum Intensity Projections to obtain a flattened image representative of the 3D volume. Time series images were processed in FIJI. The FIJI plugin StackReg (87) was used to correct for slight XY animal movements occurring during acquisition (using rigid body as correction option). Time series were finally ensembled using Kapwing Studio online tool (Kapwing Resources, San Francisco, California).

To obtain fluorescence intensity, different ROIs were drawn for cilia, PCMC, distal dendrite neuron cell body or AMsh cell body. The fluorescence of each ROI was calculated. Background fluorescence was subtracted using the following formula (CTFC = Integrated Density - (Area of selected cell X Mean fluorescence of background readings) (88). For Glia/Neuron fluorescence ratios we calculated obtaining the CTFC fluorescence values as stated above. Then the ROI’s CTFC values in AMsh was divided by CTFC values in neurons to obtain a Glia/Neuron fluorescence ratio.

To quantify the export of GCY-22-wrmScarlet from ASER. All fluorescent vesicles located within AMsh cell body were manually counted on screen across the whole AMsh volume. CFP/mEGFP channel images were used to identify the limits of the AMsh cell body.

To quantify neuron nerve receptive endings morphologies, maximum intensity projections were analyzed to quantify cilia length (measured from the cilia tip to the enlargement at cilia base), for cilia base area (excluding the cilium proper, up to the distal dendrite, based on the enriched GCY-22-wrmScarlet). ASER morphological classification is based on the number of tubulated (diameter <1 um) and non tubulated diverticula (not showing any sign of pinch) and originating from a constant 4 μm^2^ cilia base. AFD morphological classification is based on the number and length of AFD microvilli (length of microvilli in N2 animals was considered to be the baseline) and on the shape of AFD cilia base (protruding base bulges, extensions and branching).

Vesicle size was calculated using Zen software line measurement tool, a line was traced from side to side at the midline of each vesicle, obtaining an approximate value of each vesicle’s diameter. To analyze the vesicle content in the animals co-expressing of TSP-6-wrmScarlet and GCY-22-mEGFP in ASER neuron, maximum intensity projections of images of a predetermined size were analyzed to determine the number of vesicles containing TSP-6-wrmScarlet alone, GCY-22-mEGFP alone and vesicles containing both TSP-6/GCY-22, percentages were determined by dividing the number of vesicles of each group to the total number of vesicles. Vesicle size was calculated using Zen software line measurement tool, a line was traced from side to side at the maximal distance point, obtaining an approximate value of each vesicle’s diameter.

To analyze co-localization of animals co-expressing TSP-6-wrmScarlet and GCY-22-mEGFP in ASER neuron we used Imaris software Version 9 (Oxford Instruments, Zurich, Switzerland). We first established a ROI containing the cilia and their base. Pearson’s coefficient was calculated by automatic thresholding of background voxels in the selected ROI. The displayed coefficient was averaged across all the animals measured to obtain an estimate of fluorescence co-localization for both proteins.

### Chemotaxis assays

To perform NaCl chemotaxis assays, we used linear salt gradients on agar plates. 9 cm Petri dishes were used as a spatial container, the gradient was prepared as follows: first, Petri dishes were elevated using an electroporation cuvette cap to tilt the plate at an approximated 5° angle, the plate was filled with a melted 2% agar solution containing 50mM of NaCl or 5mM of CuSO_4_ (See Figure 6 – figure supplement 1A-B). Once the first layer of solution had solidified, the plate was positioned on a flat surface and a second solution of melted 2% agar (<50°C) containing 0mM of NaCl or of CuSO_4_ was poured on top. Plates were left open and the gradient was allowed to be established for approximately 1 hour prior to the assay. 30 animals were positioned in the center off-food and allowed to navigate through the plate for 30 min. At this given time point, animal’s position was annotated. To score the chemotaxis index, the plate was divided into 2 scoring regions and the chemotaxis index was calculated following the formula described in Figure 6 - figure supplement 1C.

### Optogenetic assay

The animals were grown from embryo to adulthood on OP50 supplemented or not with 200 μM of all-trans-Retinal (Sigma, #R2500) in the dark. Transgenic L4 were selected the day before and placed on fresh OP50 supplemented or not with 200 μM of all-trans-Retinal (Sigma, #R2500) the day before the assay. Blue light (480 nm, 15mw/mm2) was used to activate channelrhodopsin. We observed the reversal response (minimum 1~2 backward head swings) during the first 10 seconds of light exposure.

### Statistical analysis

Statistical analyses were performed using GraphPad Prism version 8.4.0 (San Diego, California, USA). For comparison between 2 groups with normally distributed data but unequal SD, Unpaired t test with Welch’s correction. For comparison between multiple groups (>3) normally distributed data and assuming equal standard deviations, One-way ANOVA was performed followed by a Dunnett’s post-hoc test to correct for multiple comparisons. Multiple comparisons were made between controls and the different experimental conditions, unless otherwise stated. For comparison between multiple groups (>3) normally distributed data but not assuming equal standard deviations, Brown-Forsythe ANOVA was performed followed by a Dunnett T3 post-hoc test to correct for multiple comparisons. Multiple comparisons were made between controls and the different experimental conditions. For comparison between multiple groups (>3) in non-normally distributed data, nonparametric Kruskal-Wallis ANOVA was performed followed by a Dunn’s post-hoc test for multiple comparisons. For comparison between multiple groups comparing two independent variables a Two-Way ANOVA was performed followed by a Sidak’s post-hoc test to correct for multiple comparisons. Experiment statistics appear in figure legends. Sample size is indicated in figures on top of each experimental group. In all performed tests, statistical significance threshold was set to alpha=0.05.

## Acknowledgements

Some strains were provided by the CGC, which is funded by NIH Office of Research Infrastructure Programs (P40 OD010440). We acknowledge Renaud Legouis and members of our lab for their input on the manuscript, Teresa Lobo for technical help, William Schaffer lab for AQ2335 strain. We thank J-M Vanderwinden and the ULB Imaging Facility facility (LiMiF) for imaging advice.

---

**Figure 1 – figure supplement 1:**
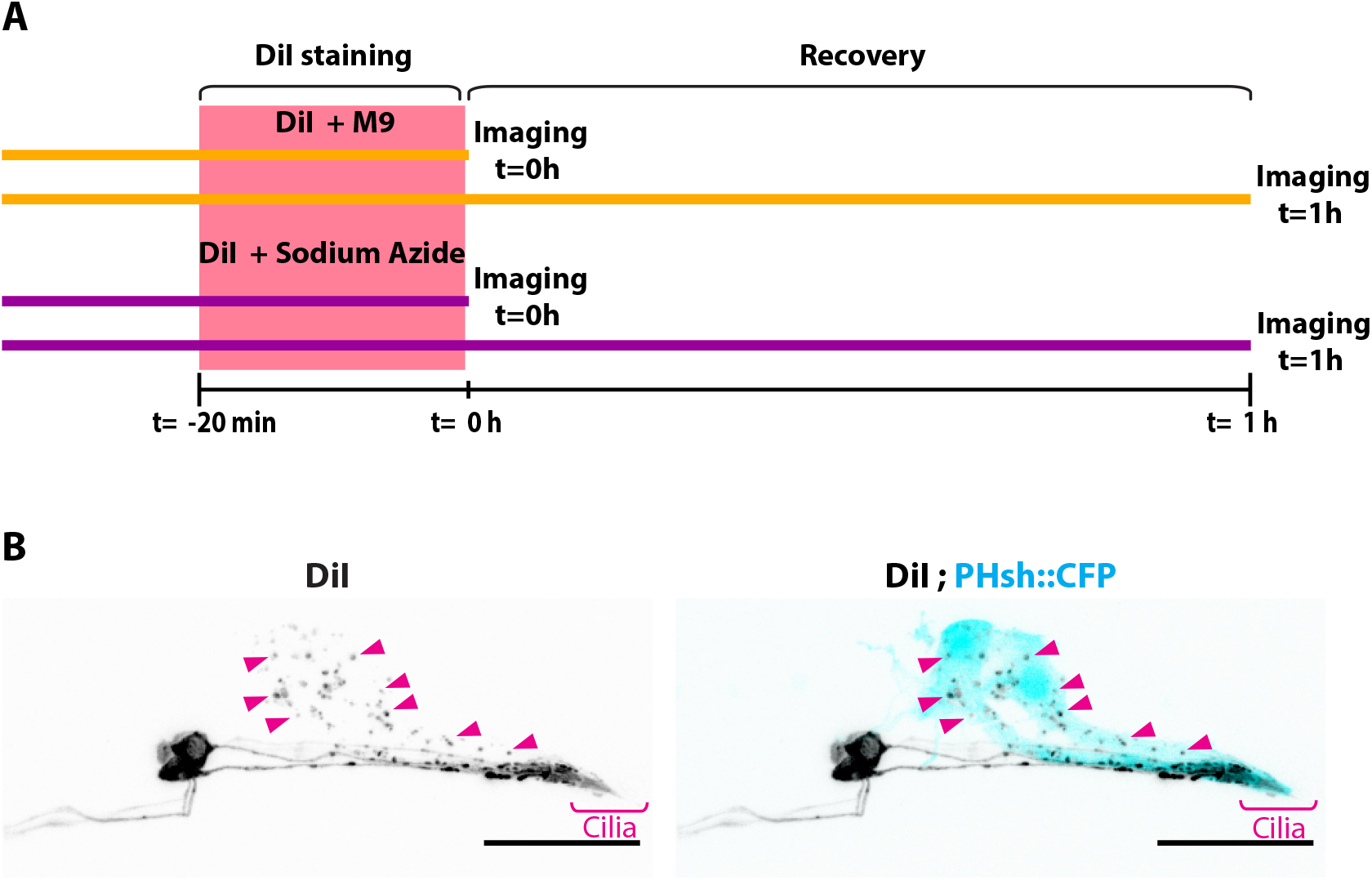
Experimental design for DiI experiment. A) Experimental scheme for Figures D-E: Animals were stained for 20 minutes with a solution of DiI in M9 buffer or with DiI in M9 buffer supplemented with 25mM Sodium Azide. The fluorescence intensity was quantified for neurons and for AMsh directly after staining (t=0) or after 1 hour of worm recovery on regular plates in absence of Sodium Azide (t=1h). B) Maximum intensity projection of DiI staining in the phasmid sensilla, PHA/B neurons show homogenous membrane staining and phasmid sheath glia contain neuronally-derived DiI vesicles (magenta arrowheads). Scale bar: 20 μm.

**Figure 2 – figure supplement 1:**
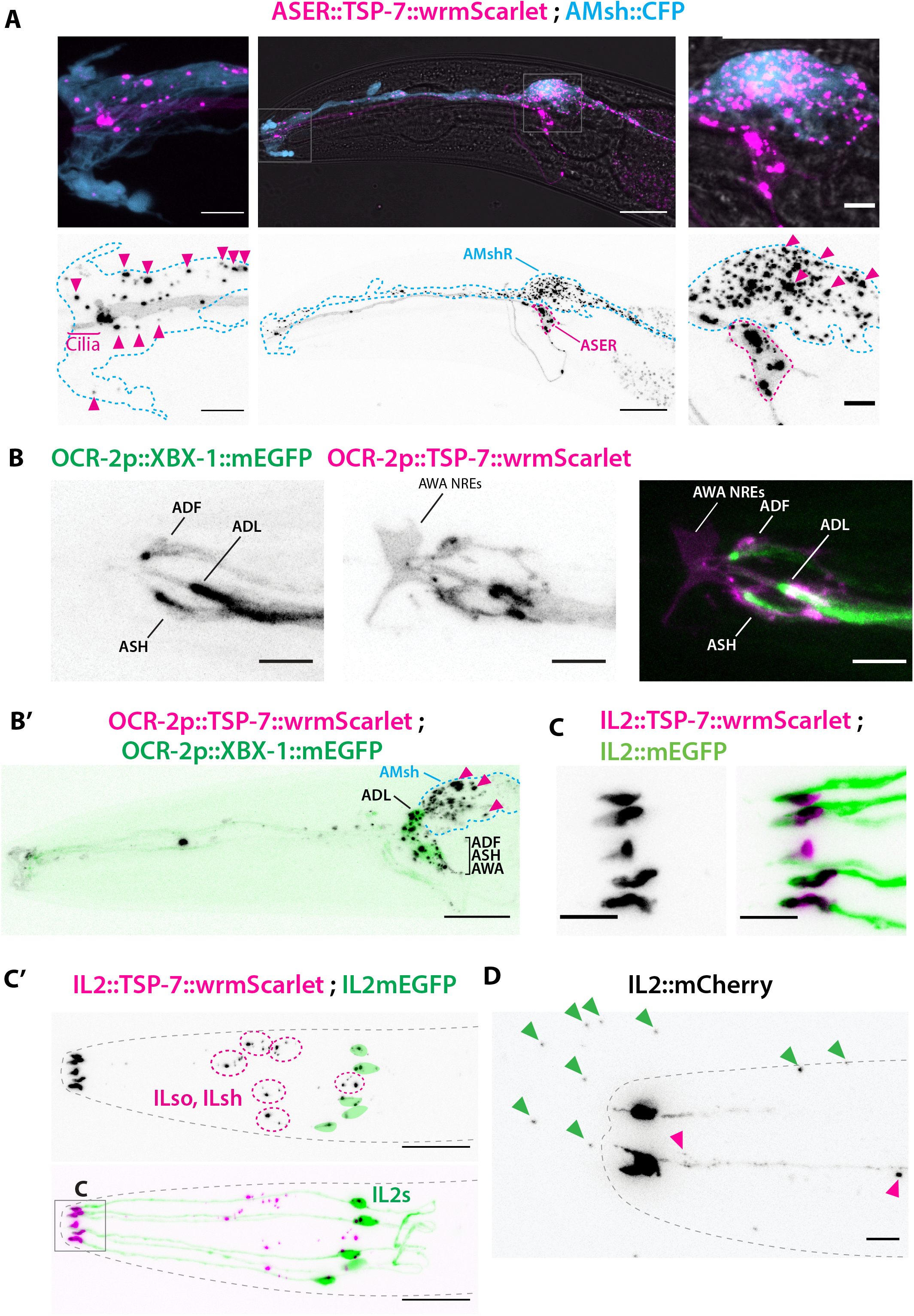
TSP-7-wrmScarlet also localizes to the cilia region, is loaded into EVs and exported to surrounding glial cells. A) Cell-specific expression of TSP-7-wrmScarlet in ASER: the fusion protein localizes to the cilia and is exported to EVs observed within the AMsh cytoplasm. Left panel shows EVs captured by AMsh glia in the surroundings of ASER cilia region (magenta arrowheads). Right panel shows TSP-7-wrmScarlet-carying EVs in the AMsh cell body (magenta arrowheads). Scale bar: 20 μm middle image and 5 μm insets. B) TSP-7-wrmScarlet expressed driven by the ocr-2 promoter (expression in ADF, ADL, ASH and AWA). TSP-7-wrmScarlet expression overlaps with the XBX-1-mEGFP marker in the cilium and PCMC of ADF, ADL, ASH and AWA. Scale bar: 5 μm. B’) In these animals, TSP-7-wrmScarlet-carrying EVs were also exported to AMsh cytoplasm (magenta arrowheads), XBX-1-mEGFP was not observed in these EVs. Theoretical position of AMsh was outlined (blue dashed line). Scale bar: 20 μm. C-C’) Co-expression of TSP-7-wrmScarlet and cytoplasmic mEGFP in IL2 neurons: C) mEGFP expression locates in all ciliary compartments while TSP-7-wrmScarlet is heavily enriched in IL2 cilia base. C’) TSP-7-wrmScarlet carrying EVs are seen within the cytoplasm of ILsh and ILso (magenta arrowheads, ILsh and ILso are outlined in magenta dashed lines). In some occasions, mEGFP and/or TSP-7-wrmScarlet-carrying EVs were also released by cilia tip into the environment (Video 3). D) Cytoplasmic mCherry expression in IL2 neurons shows EVs that have already been environmentally released (green arrowheads). Scale bar: 5 μm in C, 20 μm in C’, 5 μm in D.

**Figure 2 – figure supplement 2:**
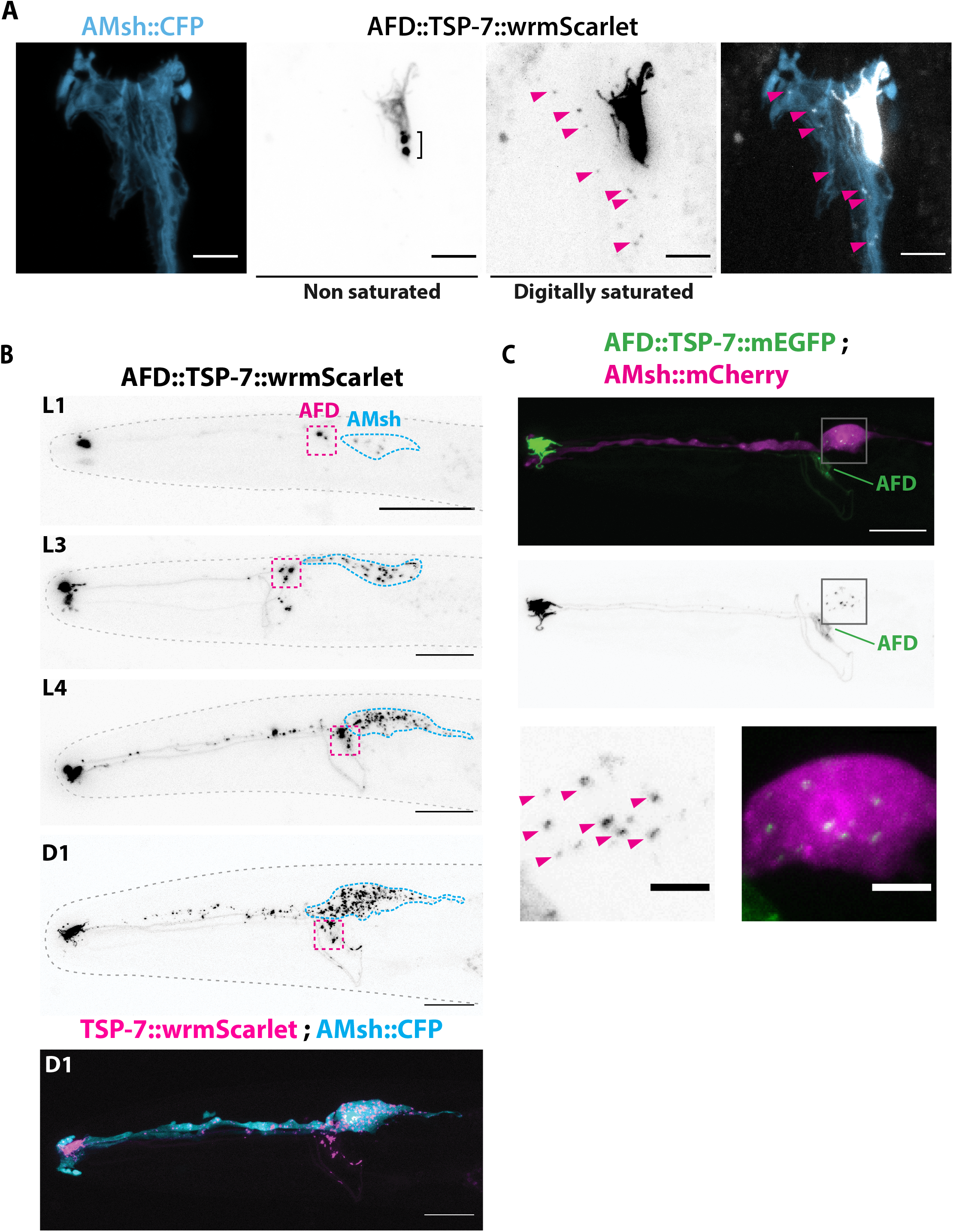
TSP-7-wrmScarlet localizes to AFD NREs, is loaded into EVs and exported to embedding AMsh glia. A) Cell-specific expression of TSP-7-wrmScarlet in AFD shows TSP-7-wrmScarlet enrichment in the microvilli and the cilia base of AFD. Non-saturated images show a TSP-7-wrmScarlet vesicular compartment within the cilia base of AFD neuron, possibly corresponding to a membrane recycling compartment. TSP-7-wrmScarlet-carrying EVs are released and captured within the cytoplasm of AMsh in the vicinity of AFD microvilli (magenta arrowheads). Image was digitally saturated by increasing the TSP-7 channel brightness in order to display the fluorescence of released vesicles. Scale bar: 5 μm. B) Longitudinal acquisitions across developmental stages from the first larval stage to Day 1 adult show the amount of TSP-7-wrmScarlet fluorescent EVs observed within AMsh cell body progressively build up as the animal ages. Scale bar: 20 μm. C) Cell-specific expression of TSP-7-mEGFP in AFD and mCherry in AMsh glia. TSP-7-mEGFP shows the same enrichment in AFD receptive endings as in A. However, TSP-7-mEGFP-carrying EVs observed in the cytoplasm of AMsh were generally reduced in number and intensity. Scale bar: 20 μm in top panels, 5 μm in bottom panels.

**Figure 3 – figure supplement 1:**
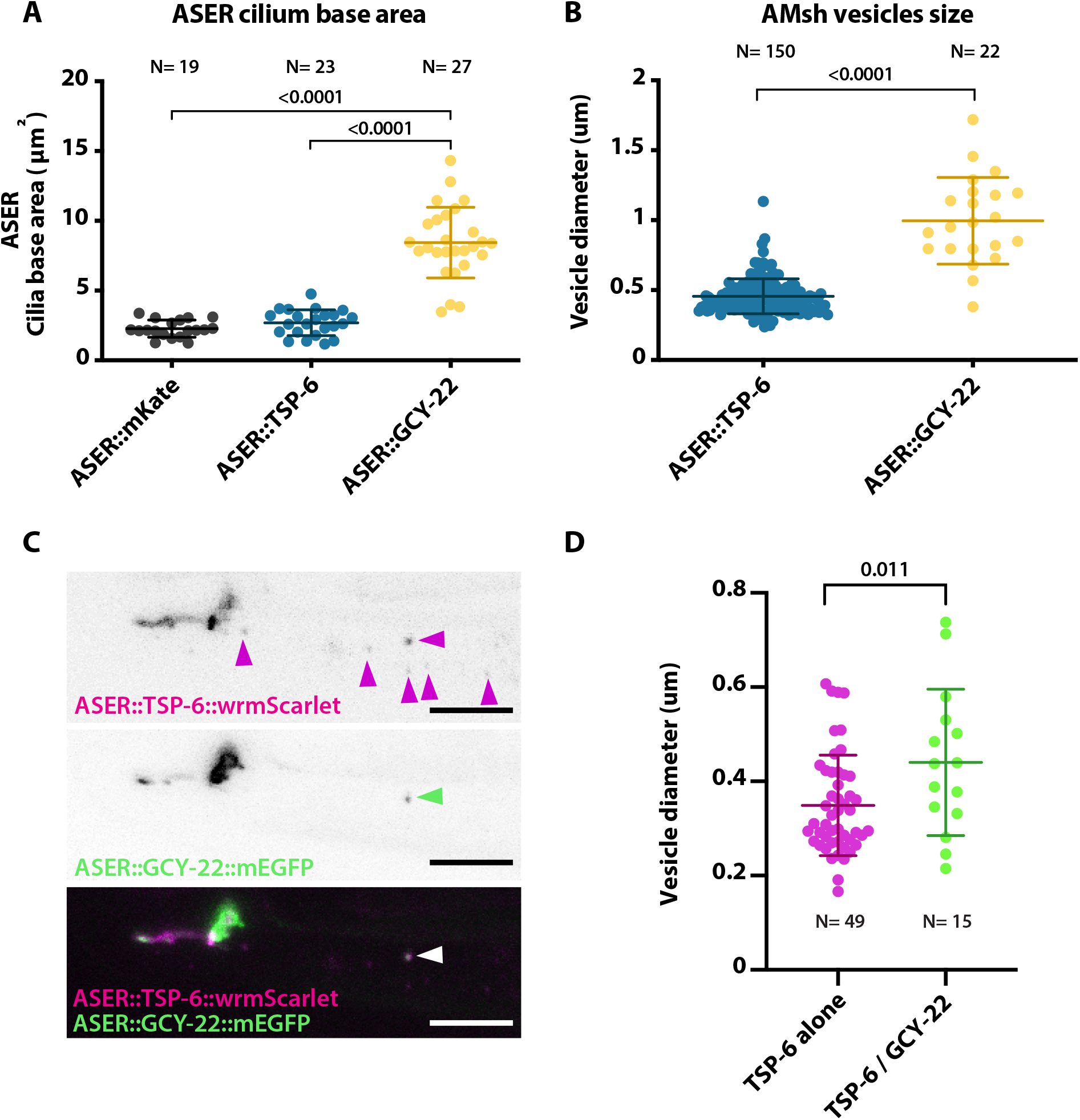
Cilia base and ectosome size is influenced by expression of TSP-6 and GCY-22. A) The PCMC area of ASER is enlarged by the expression of GCY-22-wrmScarlet compared to the expression of cytoplasmic mKate or TSP-6-wrmScarlet. One-way ANOVA, multiple comparisons corrected by Tukey test. B) The diameter of exported EVs from ASER differs between TSP-6-wrmscarlet and GCY-22-wrmscarlet-containing vesicles. Vesicles were only measured in the vicinity of ASER cilium. Unpaired t test with Welch’s correction. C) ASER cilium co-expressing TSP-6-wrmscarlet and GCY-22-mEGFP show both markers get enriched in ASER cilia, but their localisation within ASER cilia was poorly correlated (r=0.378 using Pearson’s coefficient). Most of the ASER-derived vesicles observed in AMsh cytoplasm carry TSP-6-wrmscarlet alone (magenta arrowheads), one vesicle carries TSP-6-wrmScarlet together with GCY-22-mEGFP (green arrowhead). Scale bar: 5 μm. D) Vesicle diameter differences are observed for vesicles carrying TSP-6-wrmscarlet alone or TSP-6-wrmscarlet together with GCY-22-EGFP. Vesicles were measured in the vicinity of ASER cilium. Unpaired t test with Welch’s correction.

**Figure 5 – figure supplement 1:**
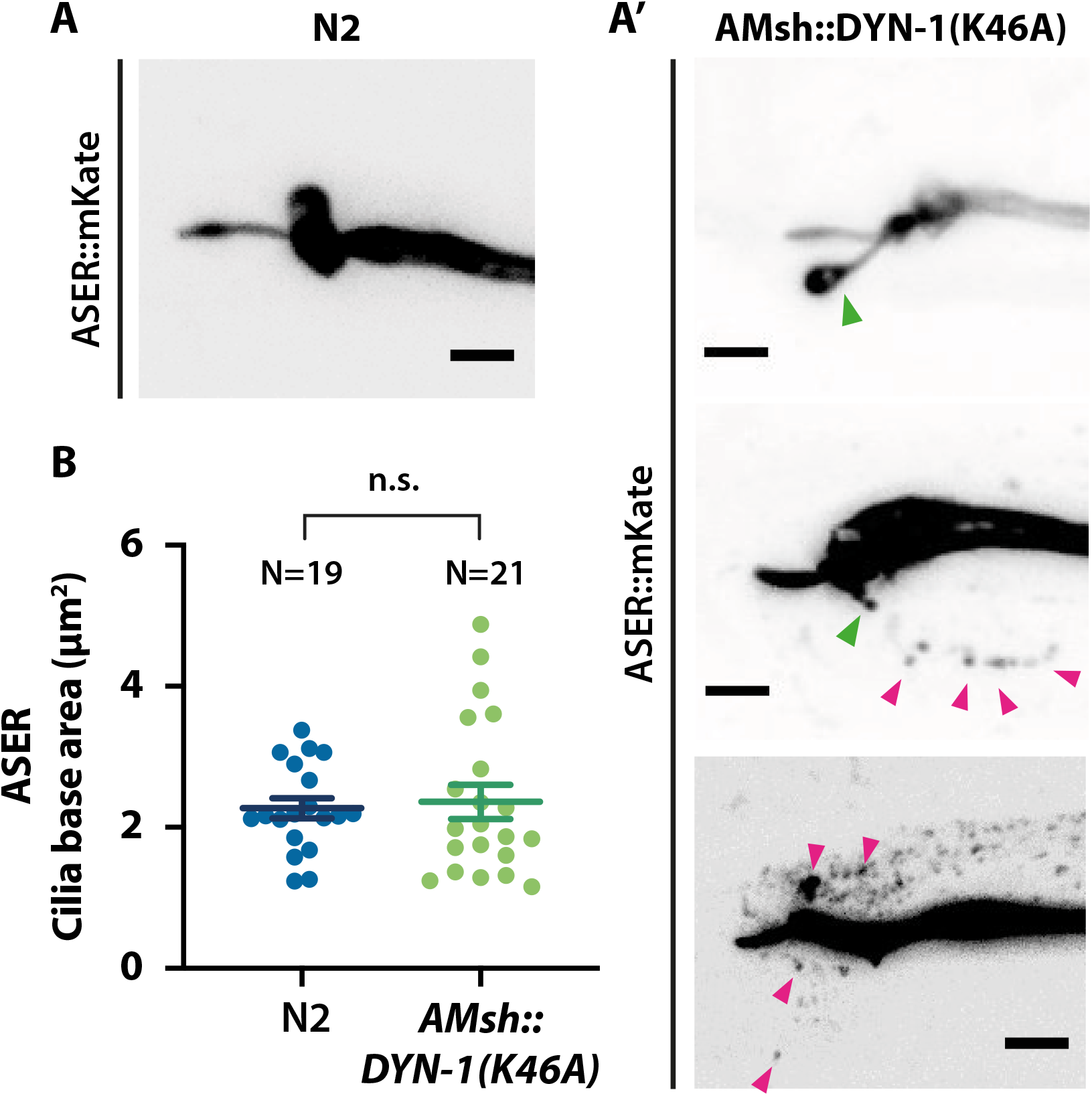
ASER sensory cilia structure is affected when AMsh phagocytic activity is disturbed. A) Cilia shape of ASER in transgenics controls expressing cytoplasmic mKate. A’) Cilia shape of ASER in transgenics expressing cytoplasmic mKate and co-expressing AMsh::DYN-1(K46A). Tubulated protrusions can be observed protruding from cilia base (green arrowheads) and the number of secreted ectosomes (magenta arrowheads) are increased in the vicinity of ASER cilia in animals expressing AMsh::DYN-1(K46A). B) The ASER cilia base area is evaluated based on 2D projections. The cilia base area is not significantly modified in AMsh::DYN-1(K46A) transgenics. Scale bar: 5 μm.

**Figure 6 – supplement 1:**
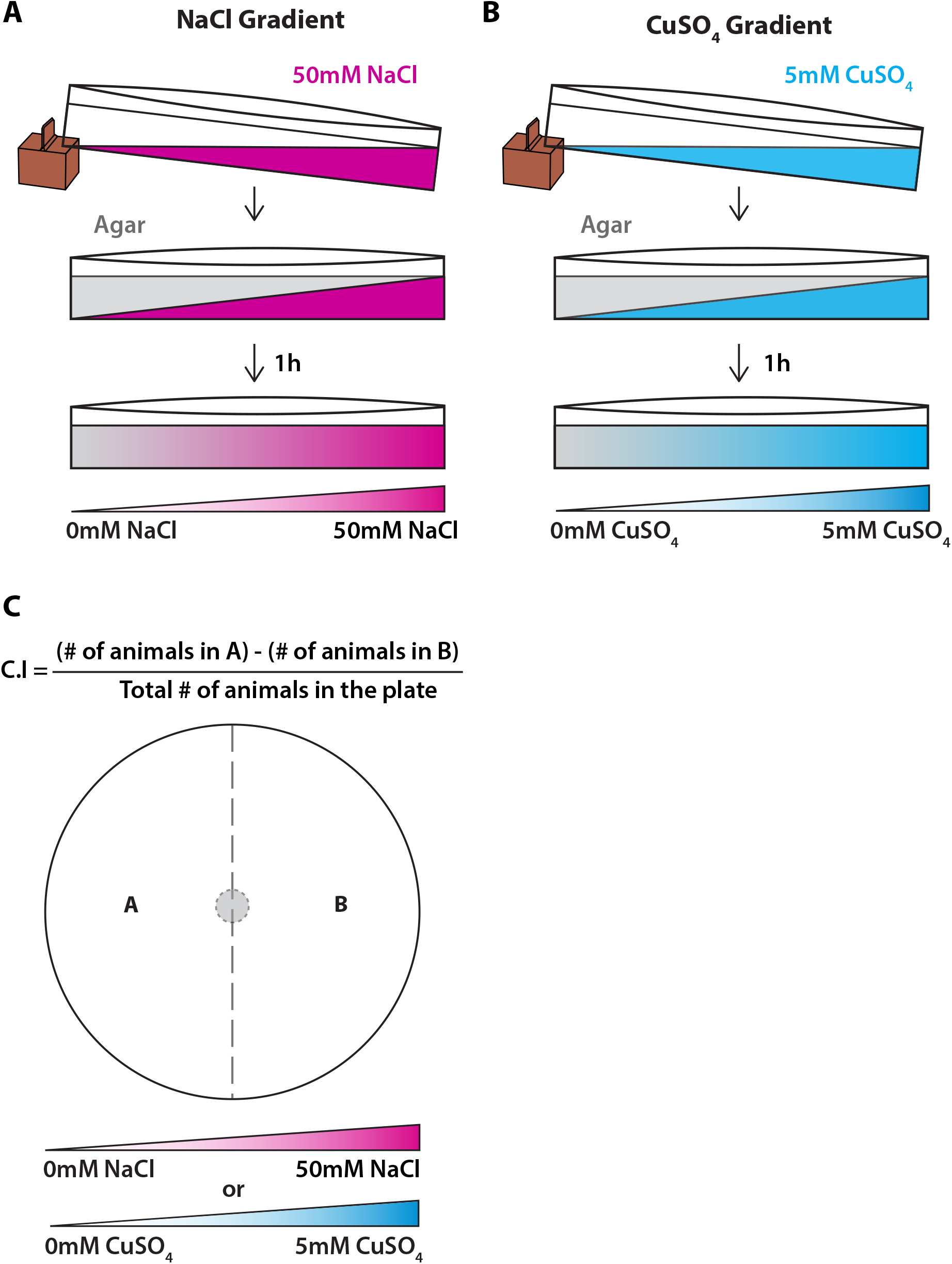
Experimental procedure followed chemotaxis gradients. A) NaCl chemotaxis gradient was generated by juxtaposition of two layers of agar solution, the bottom layer contained 50mM NaCl and the top layer did not contain NaCl, effectively creating a 5mM/cm linear gradient. B) CuSO_4_ chemotaxis gradient was generated by juxtaposition of two layers of agar solution, the bottom layer contained 5 mM CuSO_4_ and the top layer did not contain CuSO_4_, effectively creating a 0.5mM/cm linear gradient. C) Scoring method used for both chemotaxis assays. Animal position was marked on the plate when the assay finished, then chemotaxis indexes (C.I.) were scored according to the formula shown above.

**Figure 7 – figure supplement 1:**
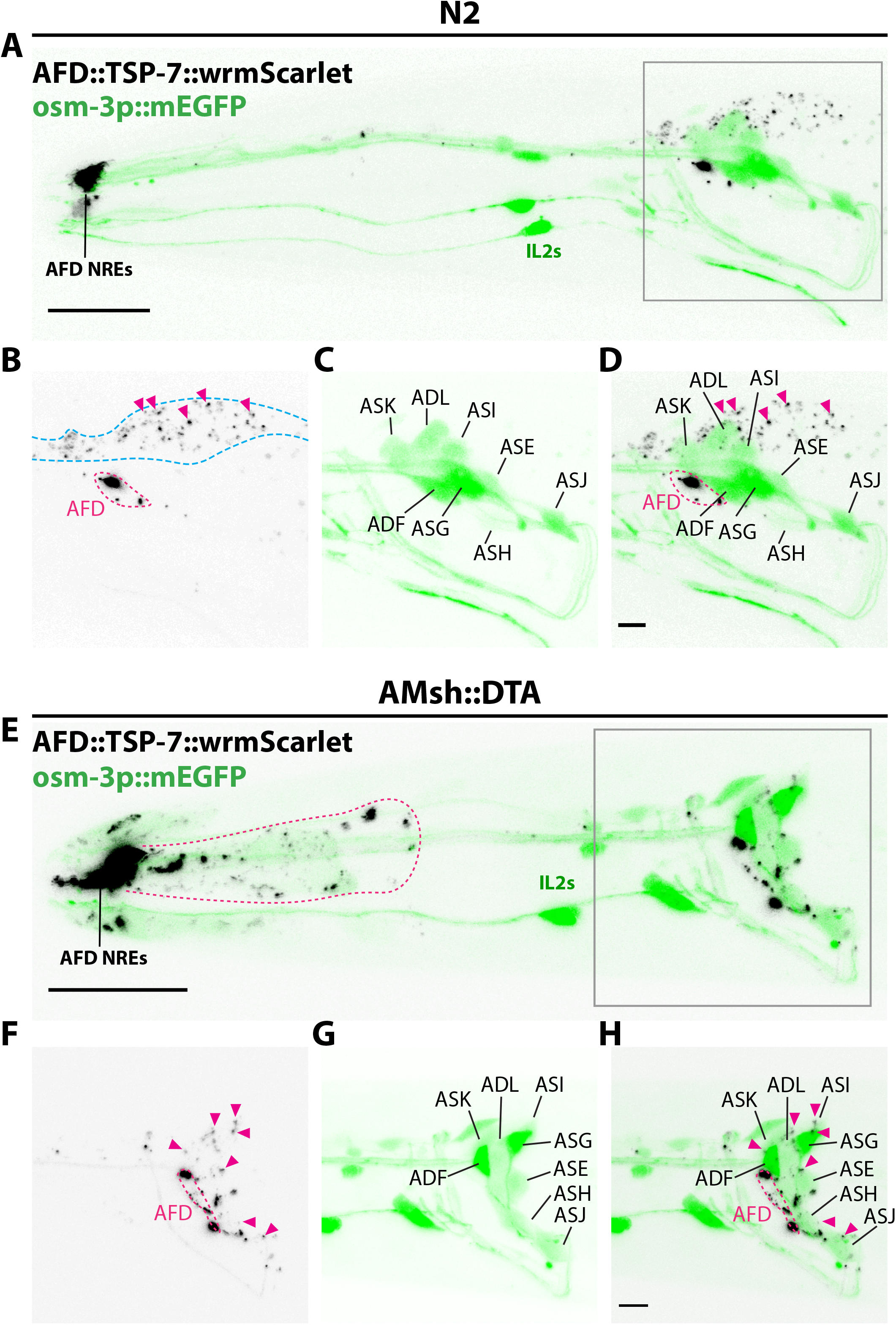
Amphid sheath ablation reroutes intake of EVs to nearby amphid neurons. A-D) Representative images of N2 animals expressing AFD::TSP-7-wrmScarlet and pOsm-3::mEGFP, driving expression in a subset of eight amphid neurons (ADF, ADL, ASE, ASG, ASH, ASI, ASJ, ASK). In wild type animals, all the EVs exported from AFD (magenta arrowheads) ends up in AMsh glia (blue dashed outline). AFD was outlined based on expression intensity of TSP-7-wrmScarlet E-H) Representative images of AMsh::DTA animals expressing AFD::TSP-7-wrmScarlet and pOsm-3::mEGFP. When glia is ablated, the EVs released from AFD (magenta arrowheads) are rerouted to nearby cells like hypoderm (magenta outline in E) and several amphid neurons. Overlap of TSP-7-wrmScarlet with GFP in the somata of the osm-3 neuronal subset allowed to confirm the presence of TSP-7-wrmScarlet EVs within ADF, ADL, ASE, ASH, ASI, ASJ (magenta arrowheads in 1F-H), although we cannot discard the presence of TSP-7-wrmScarlet in non-mEGFP labelled amphid neurons. *Scale bar: 20 μm for top head images, 5 μm for insets*.

**Video 1: EVs are released from the DiI-uptaking amphid neurons and are captured by the surrounding glia:** An animal expressing AMsh cytoplasmic CFP and stained with DiI was immobilized with 10 mM tetramisole and recorded during approximately 8 minutes. Membrane fragments become detached from ciliary regions, these DiI-carrying vesicles are sequentially trapped by glial cells (yellow arrows). Dashed arrow indicates directional flow of vesicles towards AMsh cell body. AMsh “pocket” is indicated and contains multiple already-exported vesicles. Scale bar: 5 μm.

**Video 2: Distal ectosomes are released from the cilia tip of IL2:** An animal expressing cytoplasmic mEGFP and TSP-7-wrmScarlet in IL2 neurons was immobilized with 10mM tetramisole and recorded during approximately 3 minutes. Two ectosomes grow from the cilia tip (yellow arrowheads), displayed ectosomes carry only mEGFP. Scission events occur at t=160 sec and t=167 sec. Scale bar: 5 μm.

**Video 3: Distal ectosomes are released from the cilia tip of IL2:** An animal expressing cytoplasmic mEGFP and TSP-7-wrmScarlet in IL2 was immobilized with 10mM tetramisole and recorded during approximately 3 minutes. Dynamics of a large distal ectosome (yellow arrow) growing from IL2 cilia tip carrying both mEGFP and TSP-7-wrmScarlet until the scission event occurs (at t=164 sec). Scale bar: 5 μm.

**Video 4: Basal ectosomes are released from the cilia base of IL2s:** An animal expressing cytoplasmic mEGFP and TSP-7-wrmScarlet in IL2 neurons was immobilized with 10mM tetramisole and recorded during 101 sec. Basal ectosomes (yellow arrows) are released and flow towards the IL2 sheath/socket glia cell bodies. An already released apical ectosome can be observed attached to the animal’s nose (green arrowhead). Scale bar: 5 μm.

**Video 5: Basal ectosomes are released form the cilia base of ASER:** An animal expressing TSP-6-wrmScarlet in ASER was immobilized with 10mM tetramisole and recorded for 365 sec. Small-sized ectosomes carrying TSP-6-wrmScarlet are released from ASER cilia base (yellow arrowhead), the released ectosomes are directed towards AMsh cell body (blue, CFP cytoplasmic expression, dashed arrow indicates directionality towards AMsh cell body). Scale bar: 5 μm.

**Video 6: Budding of basal ectosomes containing GCY-22-wrmScarlet originating from ASER cilium:** An animal expressing GCY-22-wrmScarlet in ASER was immobilized with 10mM tetramisole and recorded during 12 min. Large basal ectosomes (~1um dimeter) carrying GCY-22-wrmScarlet (magenta) are observed being released from the cilia base of ASER, released material ends up in AMsh cell (blue, AMsh::CFP cytoplasmic expression). GCY-22-wrmScarlet channel is shown in inverted LUT on the right. Scission events occur at t=4 sec and t=348 sec (yellow arrow indicates the start of each scission events). Scale bar: 5 μm.

**Video 7: AFD microvilli fragments are released and subsequently captured by AMsh glia:** An animal expressing TSP-6-wrmScarlet in AFD was immobilized with 10 mM tetramisole and recorded during approximately 8 min. Microvilli tips sometimes get detached from AFD NREs and are subsequently captured by AMsh (blue, AMsh::CFP cytoplasmic expression). Scission events occur at t=125 sec (yellow arrowhead indicates the start of the scission event). Scale bar: 5 μm.

