## Supplementary Tables 1-3 for "Ectosome uptake by glia sculpts *Caenorhabditis elegans* sensory cilia"

**Supplementary file 1. List of strains used in this study**

| Strain | Background allele | Description | Source |
|---|---|---|---|
| **N2** | N2 | N2 Bristol [ID=WBStrain00000001] | CGC |
| **OQ309** | N2 | Ex[pF16F9.3::CFP] | This study |
| **OQ276** | N2 | Ex[pGcy-5::TSP-6::wrmScarlet + pF16F9.3::CFP] | This study |
| **OQ319** | N2 | Ex[pSra-6::TSP-6::wrmScarlet + pF16F9.3::CFP] | This study |
| **OQ280** | N2 | Ex[pGcy-8::TSP-6::wrmScarlet + pF16F9.3::CFP] | This study |
| **OQ316** | N2 | Ex[pKlp-6::TSP-6::wrmScarlet + pF16F9.3::CFP] | This study |
| **OQ258** | N2 | Ex[pGcy-5::TSP-7::wrmScarlet + pF16F9.3::CFP] | This study |
| **OQ235** | N2 | Ex[pOcr-2::XBX-1::mEGFP + pOcr-2::TSP-7::wrmScarlet] | This study |
| **OQ205** | N2 | Ex[pKlp-6::TSP-7::wrmScarlet + pKlp-6::mEGFP::let858 3'UTR] | This study |
| **OQ157** | N2 | Ex[pKlp-6::TSP-7::wrmScarlet + pUnc-122::GFP] | This study |
| **OQ211** | N2 | Ex[pGcy-8::TSP-7::wrmScarlet + pF16F9.3::CFP] | This study |
| **OQ217** | N2 | Ex[pGcy-8::TSP-7::mEGFP + pF16F9.3::mCherry] | This study |
| **OQ200** | N2 | Ex[pGcy-8::GCY-8::wrmScarlet + pF16F9.3::CFP ] | This study |
| **OQ266** | N2 | Ex[pGcy-8::SRTX-1::wrmScarlet + pF16F9.3::CFP] | This study |
| **OQ272** | N2 | Ex[pSrbc-64::SRBC-64::wrmScarlet + pF16F9.3::CFP] | This study |
| **OQ270** | N2 | Ex[pGcy-5::GCY-22::wrmScarlet + pF16F9.3::CFP] | This study |
| **OQ325** | N2 | Ex[pGcy-5::mKate + pF16F9.3::CFP] | This study |
| **OQ322** | N2 | Ex[pGcy-5::GCY-22::mEGFP + pGcy-5::TSP-6::wrmScarlet] | This study |
| **DR86** | *daf-19(m86)* | | CGC |
| **OQ334** | *daf-19(m86)* | daf-19(m86) ; OQ270 | This study |
| **MX52** | bbs-8(nx77) | | CGC |
| **OQ311** | bbs-8(nx77) | bbs-8(nx77); OQ270 | This study |
| **GOU2366** | che-3(cas511[gfp::che-3(K2935Q)]) | | CGC |
| **OQ335** | che-3(cas511[gfp::che-3(K2935Q)]) | che-3(cas511[gfp::che-3(K2935Q)]); OQ270 | This study |
| **PR678** | tax-4(p678) | | CGC |
| **OQ315** | tax-4(p678) | tax-4(p678) ; OQ270 | This study |
| **MT9958** | ced-10(n3246) | | CGC |
| **OQ312** | ced-10(n3246) | ced-10(n3246) ; OQ270 | This study |

| | | | |
|---|---|---|---|
| **OQ303** | N2 | Ex[F16F9.3::DYN-1(K46A)::SL2mEGFP] Line 1 | This study |
| **OQ304** | N2 | Ex[F16F9.3::DYN-1(K46A)::SL2mEGFP] Line 2 | This study |
| **OQ305** | N2 | Ex[F16F9.3::DYN-1(K46A)::SL2mEGFP] Line 3 | This study |
| **OQ336** | N2 | OQ303 ; OQ270 | This study |
| **OQ327** | N2 | Ex[F16F9.3::DYN-1(K46A)::SL2mEGFP + pGcy-5::mKate] | This study |
| **OQ331** | N2 | Ex[F16F9.3::DYN-1(K46A)::SL2mEGFP + pGcy-8::mKate] | This study |
| **AQ2335** | lite-1 | lite-1; Is[psra-6Chr2-RFP] | William Schaffer Lab |
| **OQ337** | lite-1 | AQ2335 ; OQ303 | This study |
| **PR767** | ttx-1(p767) | | CGC |
| **OQ171** | ttx-1(p767) | Ex[pGcy-8::TSP-7::wrmScarlet + pF16F9.3::CFP] | This study |
| **SP1735** | dyf-7(m537) | | CGC |
| **OQ252** | dyf-7(m537) | Ex[pGcy-8::TSP-7::wrmScarlet + F16F9.3::CFP] | This study |
| **OQ231** | | Ex[pGcy-8::TSP-7::wrmScarlet + pGcy-8::mEGFP] | This study |
| **OS2248** | nsIs109 | nsIs109 [F16F9.3p::DTA(G53E) + unc-122::GFP] | Bacaj T, et al. Science. 2008 Oct 31;322(5902):744-7 |
| **OQ233** | nsIs109 | nsIs109 + Ex[pGcy-8::TSP-7::wrmScarlet + pGcy-8::mEGFP] | This study |
| **OQ338** | | Ex[pGcy-8::TSP-7::wrmScarlet + pOsm-3::mEGFP] | This study |
| **OQ340** | nsIs109 | nsIs109 + Ex[pGcy-8::TSP-7::wrmScarlet + pOsm-3::mEGFP] | This study |

**Supplementary Table 2. List of Molecular Biology / Primers used in this study**

| Gene | Fragment type | Size (in bp) | Description | Primer | Sequence |
|---|---|---|---|---|---|
| **pF16F9.3** | Promoter | 588 | Promoter for AMsh, AMso and PHsh glia | Forward | catcaaattcaacaacatgaaatg |
| | | | | Reverse | attttgtttcttactgtcttgggtatt |
| **pGcy-5** | Promoter | 1982 | Promoter for ASER neuron | Forward | tatacatgaaatacatacatagaca |
| | | | | Reverse | taatttttcgaaaacaataaatagtaaa |
| **pGcgy-8** | Promoter | 2186 | Promoter for AFD neuron | Forward | agcaaagggcgtcgattatctcgaa |
| | | | | Reverse | tttgatgtggaaaaggtagaatcgaaaatc |
| **pKlp-6** | Promoter | 1531 | Promoter for 6 pairs of IL2 neurons | Forward | caccaaaaaattcattaaagcatt |
| | | | | Reverse | cattattctgaaaagttcaactaataa |
| **pOcr-2** | Promoter | 2500 | Promoter for AWA, ASH, ADL and ADF neurons | Forward | ttgtacagtttacatttattataggtaggcacta |
| | | | | Reverse | cttaatgatgtgatgtactctactgataaga |
| **pSrbc-64** | Promoter | 1807 | Promoter for ASK neuron | Forward | gttttctaaaaatgagatattactagtg |
| | | | | Reverse | cagactgtgacaagaaaactgaa |
| **pOsm-3** | Promoter | 2511 | Promoter for ASH, ASI and PVQ neurons | Forward | cgaggtttgcttcaaaattcggta |
| | | | | Reverse | tccgacgcatagctggaaat |
| **CFP** | Fluorescent protein | 870 | Cytoplasmic expression of CFP | Forward | atgagtaaaggagaagaacttttcac |
| | | | | Reverse | ctatttgtatagttcatccatgcc |
| **mEGFP** | Fluorescent protein | 872 | Cytoplasmic expression of mEGFP | Forward | atgtccaagggagaggagctct |
| | | | | Reverse | ctacttgtagagctcgtccattccg |
| **mKate** | Fluorescent protein | 849 | Cytoplasmic expression of mKate | Forward | atgtccgagctcatcaaggag |
| | | | | Reverse | ttaacggtgtccgagcttgg |
| **mCherry** | Fluorescent protein | 864 | Cytoplasmic expression of mKate | Forward | atggtctcaaagggtgaagaagataa |
| | | | | Reverse | ttacttatacaattcatccatgccacct |
| **tsp-6** | Gene | 1577 | From genomic sequence | Forward | atggttcaaggatgtggtaacaaatgc |
| | | | | Reverse | agcttgggagcgtttctctttg |
| **tsp-7** | Gene | 2952 | From genomic sequence | Forward | atggtagaaggaggagttaccatagtta |
| | | | | Reverse | ataataaaagtcatggaaatccttgaga |
| **gcy-22** | Gene | 4621 | From genomic sequence | Forward | atgagtttcatatcaaaatgttttatttgc |
| | | | | Reverse | gatagattctccattctccttcgc |
| **xbx-1** | Gene | 2270 | From genomic sequence | Forward | atgaacatttgggatcttgcca |
| | | | | Reverse | tcgaacattaatttttgcgattcgat |
| **gcy-8** | Gene | 5431 | From genomic sequence | Forward | atgcgaacaaagaaggcattt |
| | | | | Reverse | tctctgcaatcctgttggatt |
| **srtx-1** | Gene | 1299 | From genomic sequence | Forward | atgttggaagatctcttgtacgaagtac |
| | | | | Reverse | ttcttgatagtagaagctgacagatcg |
| **srbc-64** | Gene | 2163 | | Forward | atgcctgaaatagtaataatcttgaaca |

| | | | | | |
|---|---|---|---|---|---|
| | | | From genomic sequence | Reverse | ctgtgaccatgtgagcacag |
| **dyn-1** | Gene | 3642 | From genomic sequence | Forward | atgtcgtggcaaaaccaggg |
| | | | | Reverse | ttatctaggcggtgccatgttg |
| **dyn-1(K46A)** | Gene | - | K46A substitution mutagenesis | Forward | atcgccgtcgtcggaggacagtccgctggagcgtcgtcggtgc |
| | | | | Reverse | tgtcctccgacgacggcgatctgtggaagttcgaagctgac |
| **wrmScarlet C-terminal Fusion** | Fluorescent protein | 699 | C-terminal fusion wrmScarlet | Forward | atggtcagcaagggagaggcagtta |
| | | | | Reverse | cttgtagagctcgtccattcctccg |
| **mEGFP C-terminal Fusion** | Fluorescent protein | 872 | C-terminal fusion mEGFP | Forward | atgtccaagggagaggagctcttca |
| | | | | Reverse | ctacttgtagagctcgtccattccg |
| **SL2-mEGFP** | Fluorescent protein | 1070 | SL2-mEGFP reporter | Forward | ccgctgtctcatcctactttcac |
| | | | | Reverse | ctacttgtagagctcgtccattccg |
| **let-858 3'UTR** | Regulatory element | 408 | let-858 3'UTR Regulatory element | Forward | attttcaaattttaaatactgaatatttgt |
| | | | | Reverse | ccaagcgaggacaattct |

**Supplementary Table 3. List of plasmids used in this study**

| Plasmid name | Description | Cloning method |
|---|---|---|
| **AS1** | pF16F9.3::CFP::let-858 3'UTR | Gateway cloning |
| **AS2** | pGcy-5::TSP-6::wrmScarlet | Gateway cloning |
| **AS3** | pSra-6::TSP-6::wrmScarlet | Gateway cloning |
| **AS4** | pGcy-8::TSP-6::wrmScarlet | Gateway cloning |
| **AS5** | pKlp-6::TSP-6::wrmScarlet | Gateway cloning |
| **AS6** | pKlp-6::mEGFP::let-858 3'UTR | Gateway cloning |
| **AS7** | pGcy-5::TSP-7::wrmScarlet | Gateway cloning |
| **AS8** | pOcr-2::XBX-1::mEGFP | Gateway cloning |
| **AS9** | pOcr-2::TSP-7::wrmScarlet | Gateway cloning |
| **AS10** | pKlp-6::TSP-7::wrmScarlet | Gateway cloning |
| **AS11** | pF16F9.3::mCherry::unc-54 3'UTR | Gateway cloning |
| **AS12** | pGcy-8::TSP-7::wrmScarlet | Gateway cloning |
| **AS13** | pGcy-8::GCY-8::wrmScarlet | Gateway cloning |
| **AS14** | pGcy-8::SRTX-1::wrmScarlet | Gateway cloning |
| **AS15** | pSrbc-64::SRBC-64::wrmScarlet | Gateway cloning |
| **AS16** | pGcy-5::GCY-22::wrmScarlet | Gateway cloning |
| **AS17** | pGcy-5::GCY-22::mEGFP | Gateway cloning |
| **AS18** | pF16F9.3::DYN-1(K46A)::SL2-mEGFP | Gateway cloning |
| **AS19** | pGcy-5::mKate::let-858 3'UTR | Gateway cloning |
| **AS20** | pGcy-8::mKate::let-858 3'UTR | Gateway cloning |
| **AS21** | pGcy-8::mEGFP::let-858 3'UTR | Gateway cloning |
| **AS22** | pOsm-3::mEGFP::let-858 3'UTR | Gateway cloning |
